# Structures of LIG1 active site mutants reveal the importance of DNA end rigidity for mismatch discrimination

**DOI:** 10.1101/2023.03.21.533718

**Authors:** Mitchell Gulkis, Qun Tang, Matthew Petrides, Melike Çağlayan

**Author notes:** Authors contributed equally (Mitchell Gulkis, Qun Tang).

## Abstract

ATP-dependent DNA ligases catalyze phosphodiester bond formation in the conserved three-step chemical reaction of nick sealing. Human DNA ligase I (LIG1) finalizes almost all DNA repair pathways following DNA polymerase-mediated nucleotide insertion. We previously reported that LIG1 discriminates mismatches depending on the architecture of the 3’-terminus at a nick, however the contribution of conserved active site residues to faithful ligation remains unknown. Here, we comprehensively dissect the nick DNA substrate specificity of LIG1 active site mutants carrying Ala(A) and Leu(L) substitutions at Phe(F)635 and Phe(F)F872 residues and show completely abolished ligation of nick DNA substrates with all 12 non-canonical mismatches. LIG1^EE/AA^ structures of F635A and F872A mutants in complex with nick DNA containing A:C and G:T mismatches demonstrate the importance of DNA end rigidity, as well as uncover a shift in a flexible loop near 5’-end of the nick, which causes an increased barrier to adenylate transfer from LIG1 to the 5’-end of the nick. Furthermore, LIG1^EE/AA^/8oxoG:A structures of both mutants demonstrated that F635 and F872 play critical roles during steps 1 or 2 of the ligation reaction depending on the position of the active site residue near the DNA ends. Overall, our study contributes towards a better understanding of the substrate discrimination mechanism of LIG1 against mutagenic repair intermediates with mismatched or damaged ends and reveals the importance of conserved ligase active site residues to maintain ligation fidelity.

## Introduction

DNA strand breaks, which can be generated because of normal DNA transactions or DNA lesion removal by base excision repair, are potentially toxic intermediates in genomic DNA, which upon DNA replication, unrepaired strand breaks can cause replication fork collapse leading to cytotoxic double strand breaks (1–3). DNA ligases are a large family of proteins responsible for catalyzing nucleophilic attack between the adjacent 3’-hydroxyl (OH) and 5’-phosphate (P) groups in nick DNA using NAD^+^ or ATP as a nucleotide cofactor to form a covalent enzyme-nucleoside monophosphate in a wide range of DNA transactions including DNA repair, replication, and recombination in all three kingdoms of life (4–6). ATP-dependent eukaryotic DNA ligases that are related in sequence and structure catalyze nick sealing through three conserved chemical and consecutive steps (4). In the first step of the ligation reaction, the ligase attacks the α-phosphate of ATP which results in an adenylate (AMP) moiety being covalently linked to the ε-amino group of an active site lysine residue resulting in the formation of the initial ligase-AMP intermediate (6,7). The adenylated ligase then undergoes a conformational change which allows the enzyme to surveille the genome until a nick is encountered. In the second step of the ligation reaction, the ligase catalyzes the transfer of AMP to the 5’-P of the nick forming a 5’-5’ phosphoanhydride high energy bond referred to as the DNA-AMP intermediate (8,9). Finally, in the third step, the ligase catalyzes in-line nucleophilic attack of the 3’-OH group onto the 5’-phosphorylated DNA end creating a phosphodiester bond and displacing AMP (10). Human DNA ligases also require a catalytic magnesium (Mg^2+^) ion for efficient catalysis during the initial attack on ATP and the nucleophilic attack on the phosphoanhydride bond (11).

DNA ligases encoded by the human *LIG1*, *LIG3*, and *LIG4* genes belong to the nucleotidyl transferase family and share a highly conserved region that compromises the catalytic core containing the oligonucleotide binding (OBD) and the adenylation (AdD) domains (1–5). The DNA binding (DBD) domain is less conserved among eukaryotic ligases and binds more robustly to nick DNA (6–10). Although they share a common catalytic core, and the ligation reaction is universally conserved, human DNA ligases exhibit distinct ligation fidelities. Among them, DNA ligase I (LIG1) is responsible for the ligation of over 10 million Okazaki fragments during each round of DNA replication while interacting with proliferating cell nuclear antigen (PCNA) and seals the nick intermediate following DNA polymerase-mediated nucleotide insertion at the final step of almost all DNA repair pathways (12,13). The expression of LIG1 is frequently altered in many types of cancer, which has been considered an indicator of altered DNA repair capacity in cells (14–17). LIG1 deficiency has been associated with mutations in the *LIG1* gene resulting in LIG1-deficiency syndrome which clinically manifests as immunodeficiency, enhanced hypersensitivity to DNA damaging agents, genomic instability, and cancer predisposition (18–20). Therefore, it’s important to understand the molecular mechanism of the ligation reaction by LIG1 to underpin how ligation fidelity is maintained during nuclear replication and DNA repair in normal *versus* disease states (21).

The first structure of LIG1 in complex with unligatable nick DNA revealed a three-dimensional catalytic mechanism of the ligation reaction involving the formation of two covalent intermediates as well as important insight into ligase fidelity (22). In this LIG1 structure, it was reported that all three domains of the ligase make close contacts with the nick DNA duplex which undergo a conformational change between the OBD and AdD domains of the protein which is required to proceed from step 1, where the initial LIG1-AMP intermediate is formed, to step 2, where the ligase engages with nick DNA to form the DNA-AMP intermediate. In addition to the active site lysine residue (K568) that is covalently attached to the AMP moiety, the roles of other LIG1 active site residues, particularly phenylalanine (Phe) residues Phe(F)635 and Phe(F)872 residing in the AdD and OBD domains, respectively, have been suggested to play critical roles for catalysis through their interactions with the DNA ends (22). The first structure of LIG1 also demonstrated that F635 and F872 are important for stabilizing the A-form conformation of the nick and for positioning DNA ends by pressing against the sugar moiety to enforce proper geometric constraints for in-line nucleophilic attack (22). Furthermore, the amino acid sequence alignment of DNA ligases shows the conservation of F635 back to *Saccharomyces cerevisiae* and the conservation of F872 back to *Escherichia coli* (Supplementary Scheme 1). Although they are highly conserved amongst ATP-dependent DNA ligases from human and other sources, the role of F635 and F872 active site residues for non-canonical substrate discrimination of a human DNA ligase at atomic resolution remains unknown.

Later solved LIG1 structures have revealed that the amino acid residues Glu(E)346 and Glu(E)592 are the high-fidelity Mg^2+^ binding sites that are independent of the catalytic Mg^2+^, which is required during the initial and final steps of the ligation reaction (23–25). Moreover, the mutations at these two glutamic acids residues (E346A/E592A or EE/AA) leads to a lower fidelity LIG1 mutant (referred to as LIG1^EE/AA^ in the present study) and enables the ligase to better accommodate a nick DNA with oxidative damage (8oxoG) at the 3’-end (23). Furthermore, our recent structures of LIG1/nick DNA complexes uncovered the ligase strategies that favor or deter ligation of A:C and G:T mismatches (26). We reported that the ligase active site can accommodate a G:T mismatch, which exhibits a similar conformation as a canonical A:T base pair during step 2 of the ligation reaction, when the DNA-AMP intermediate is formed, while it stays in the initial LIG1-AMP intermediate state, during step 1, in the presence of an A:C mismatch at the 3’-end (26). Although previous findings reported the mechanism by which LIG1 surveils DNA ends during nick sealing (22–26), it is entirely missing at atomic resolution how the ligase active site contributes to ligation fidelity while LIG1 is engaging with mutagenic repair intermediates with damaged or mismatched DNA ends.

In the present study, we generated Phe(F) to Ala(A) or Leu(L) mutations at F635 and F872 in the wild-type and low-fidelity (LIG1^EE/AA^) background of LIG1, and comprehensively characterized the single (F635A, F635L, F872A, F872L) and triple (LIG1^EE/AA^ F635A, LIG1^EE/AA^ F872A) mutants at biochemical and structural levels. We investigated the substrate specificity of these LIG1 mutants in ligation assays *in vitro* and demonstrated completely abolished nick sealing for all 12 non-canonical mismatches. Additionally, we solved the structures of LIG1^EE/AA^ F635A and F872A mutants with nick DNA complexes containing a cognate A:T, mismatches A:C and G:T, and oxidatively damaged 8oxoG:A ends at the 3’-end. Our mismatch structures for both LIG1^EE/AA^ mutants revealed conformational changes at the DNA ends, as well as a shift in the flexible loop between R736 and R741, which results in the putative formation of additional hydrogen bonds between R738 and the 5’-P of the nick during step 1 of the ligation reaction, which together hinders the ligase for further catalytic steps, in contrast to the LIG1^EE/AA^/A:T structures which were captured during step 2 when the DNA-AMP intermediate is formed. Furthermore, our LIG1^EE/AA^ F872A/8oxoG:A structure demonstrated a similar DNA end misalignment and the R738 loop shift during the initial step of the ligation reaction and a lack of ligation *in vitro*, while we captured LIG1^EE/AA^ F635A/8oxoG:A during step 2 and observed the mutagenic nick sealing of damaged end. Overall, our study contributes to elucidating the mechanism by which LIG1 discriminates against mutagenic repair intermediates with mismatched or damaged ends and the contribution of conserved F635 and F872 residues to faithful ligation during nuclear replication and DNA repair.

## Methods

### Preparation of LIG1 active site mutant constructs

We generated Phe(F) to Ala(A) and Leu(L) amino acid substitutions at F635 and F872, residues that reside in Adenylation domain (AdD) and Oligonucleotide-binding domain (OBD) of DNA ligase I (LIG1), respectively, within the catalytic core of the enzyme (Supplementary Scheme 2). Plasmid DNA constructs carrying single mutations F635A, F635L, F872A, F872L, double-mutation E346A/E592A (LIG1^EE/AA^), and triple mutations E346A/E592A/F635A (LIG1^EE/AA^ F635A) and E346A/E592A/F872A (LIG1^EE/AA^ F872A) were prepared using the wild-type LIG1 C-terminal (△261) construct as reported previously (26–33). Plasmid DNA constructs were cloned into pET-24b (Novagen) expression vector by site-directed mutagenesis and the coding sequences of all mutants were confirmed by DNA sequencing prior to use.

### Preparation of nick DNA substrates for biochemical and structural studies

Nick DNA substrates with a 6-carboxyfluorescein (FAM) label at 3’-end were used in DNA ligation assays. Nick DNA substrates containing all 12 mismatches were prepared by annealing oligonucleotides 3’-dA, dT, dG, or dC with template base A, T, G, or C (Supplementary Table 1). Nick DNA substrates containing 8-oxodG were prepared by annealing 3’-8oxodG with template base A or C (Supplementary Table 2). Nick DNA substrates containing A:T, A:C, G:T, and 8oxoG:A were used in the LIG1 X-ray crystallization experiments (Supplementary Table 3).

### Protein purifications

Human his-tag LIG1 C-terminal (△261) wild-type, single mutants F635A, F635L, F872A, and F872L, double mutant LIG1^EE/AA^, and triple mutants LIG1^EE/AA^ F635A and F872A were purified as described previously (26–33). Briefly, the proteins were overexpressed in *E. coli* Rosetta (DE3) pLysS cells and grown in Terrific Broth (TB) media with kanamycin (50 μgml^−1^) and chloramphenicol (34 μgml^−1^) at 37 °C. Once the OD_600_ reached 1.0, cells were induced with 0.5 mM isopropyl β-D-thiogalactoside (IPTG) and overexpression continued overnight at 20 °C. After centrifugation, cells were lysed in lysis buffer containing 50 mM Tris-HCl (pH 7.0), 500 mM NaCl, 20 mM imidazole, 2 mM β-mercaptoethanol, 5% glycerol, and 1 mM PMSF by sonication at 4 °C. The lysate was pelleted at 31,000 x g for 90 min at 4 °C. The cell lysis solution was clarified and then loaded onto a HisTrap HP column that was previously equilibrated with binding buffer containing 50 mM Tris-HCl (pH 7.0), 500 mM NaCl, 20 mM imidazole, 2 mM β-mercaptoethanol, and 5% glycerol. The column was washed with binding buffer and then eluted with an increasing imidazole gradient 0-500 mM at 4 °C. The collected fractions were then subsequently loaded onto a HiTrap Heparin column that was equilibrated with binding buffer containing 20 mM Tris-HCl (pH 7.0), 50 mM NaCl, 2 mM β-mercaptoethanol, and 5% glycerol. The protein was eluted with a linear gradient of NaCl up to 1 M. LIG1 proteins were further purified by Superdex 200 10/300 column in the buffer containing 50 mM Tris-HCl (pH 7.0), 200 mM NaCl, and 1 mM DTT, and 5% glycerol. LIG1^EE/AA^ F635A and F872A proteins which were used for crystallization experiments were exchanged to crystal storage buffer containing 20 mM Tris-HCl (pH 7.0), 200 mM NaCl, and 1 mM DTT. All proteins purified in this study were concentrated, aliquoted, and stored at −80 °C. Protein quality was evaluated on a 10% SDS-PAGE gel, and protein concentrations were measured using absorbance at 280 nm.

### DNA ligation assays

Ligation assays were performed to investigate the substrate specificity of LIG1 wild-type, LIG1^EE/AA^, and six active site mutants (Supplementary Scheme 3). For this purpose, we used the nick DNA substrates including all 12 non-canonical 3’-preinserted mismatches as well as 3’-preinserted 8-oxoG:A and 8oxoG:C (Supplementary Tables 1 and 2). The reaction was initiated by the addition of LIG1 (150 nM) to a mixture containing 50 mM Tris-HCl (pH 7.5), 100 mM KCl, 10 mM MgCl_2_, 1 mM ATP, 1 mM DTT, 100 µgml^−1^ BSA, 10% glycerol, and the nick DNA substrate (500 nM) in a final volume of 10 µl. The reaction mixture was incubated at 37 °C and stopped at the time points indicated in the figure legends by mixing with an equal volume of loading dye containing 95% formamide, 20 mM ethylenediaminetetraacetic acid, 0.02% bromophenol blue, and 0.02% xylene cyanol. Reaction products were separated by electrophoresis on an 18% Urea-PAGE gel, the gels were scanned with Typhoon PhosphorImager RGB (Amersham), and the data was analyzed using ImageQuant software as described previously (26–33). DNA ligation reactions were performed similarly for wild-type and all mutants of LIG1.

### Fluorescence-based thermal shift assays

Thermal shift assays were carried out in 96 well plates using the CFX96 RT-PCR detection system (Bio-Rad). Thermal stability experiments were performed in the reaction mixture containing LIG1 (5 µM) in the absence or presence of nick DNA (50 µM) in a total volume of 25 µl. Nick DNA substrate including preinserted 3’-dG:C was used (Supplementary Table 4). After incubation at 4 °C for 1 h, 20X Sypro Orange protein dye (Invitrogen) was added, and the plate was then sealed with an optical seal and centrifuged. The thermal scan ranged from 10-95 °C with a temperature ramp rate of 0.20 °C/min. The fluorescence intensity upon binding of Sypro Orange was measured with excitation/emission of 533/580 nm. Data analysis and report generation were performed by the Maestro instrument software (Biorad). T_m_ values were calculated manually from the negative derivative plot at the point of inflection of the curve (the midpoint for protein unfolding). All images were drawn using Graph Pad Prism 9. Thermal shift assays were performed in at least biological duplicates each containing technical triplicates for LIG1 wild-type and all mutants tested in this study.

### Crystallization and structure determination

LIG1^EE/AA^ C-terminal (△261) F635A and F872A mutants were used for the crystallization experiments. The EE/AA mutation aids in crystallization (23–26). All LIG1-DNA complex crystals were grown at 20 °C using the hanging drop method as described previously (26). LIG1 (at 26 mgml^−1^)/DNA complex solution was prepared in 20 mM Tris-HCl (pH 7.0), 200 mM NaCl, 1 mM DTT, 8 mM EDTA, and 4 mM ATP at 1.5:1 DNA:protein molar ratio and incubated at 37 °C for 3 min. 1 μl of DNA:protein complex was mixed with 1 μl reservoir solution to form the hanging drop (Supplementary Table 5). All crystals grew in 1-2 days. Crystals were harvested and submerged in cryoprotectant solution containing reservoir solution mixed with glycerol to a final concentration of 20% glycerol before being flash cooled in liquid nitrogen for data collection. Crystals were kept at 100 °K during X-ray diffraction data collection using the beamline 22-ID (APS). Diffraction images were indexed and integrated using HKL2000 (HKL Research, Inc). All structures were solved through molecular replacement using PHASER with PDB entry 7SUM as the search model (34). Iterative rounds of model building in COOT and refinement with PHENIX or REFMAC5 were used to produce the final models (35–37). 3DNA was used for sugar pucker analysis (57). All structural images were drawn using PyMOL (The PyMOL Molecular Graphics System, V0.99, Schrödinger, LLC). Detailed crystallographic statistics are provided in Table 1.

**Table 1.**
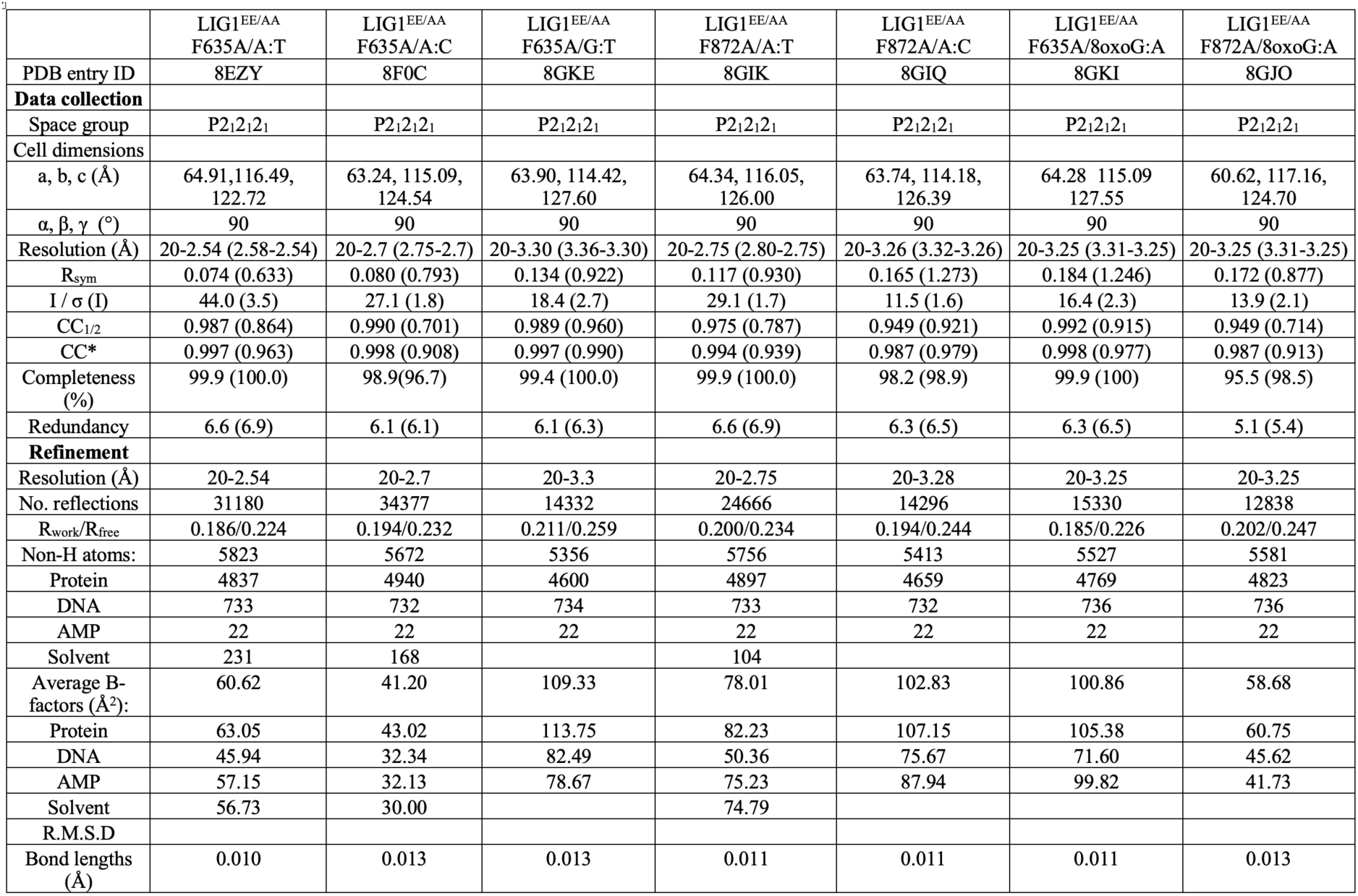
X-ray data collection and refinement statistics of LIG1^EE/AA^ F635A and F872A in complex with nick DNA duplexes with A:T, A:C, G:T, and 8oxoG:A at the 3’-end.

## Results

### Structures of LIG1 reveal the roles of F635 and F872A active site residues for mismatch discrimination

In the structure of LIG1^EE/AA^/nick DNA containing a cognate A:T base pair (26), the ligase active site residues F635 and F872 insert into the minor groove where the aromatic moiety presses against the deoxyribose sugar at the 3’- and 5’-ends of the nick DNA, respectively (Figure 1A-C). To elucidate the importance of these conserved residues to ligation fidelity at atomic resolution, we generated F635A and F872A mutations in the low-fidelity background of LIG1 (EE/AA) and solved the X-ray structures of LIG1^EE/AA^ F635A and F872A mutants with nick DNA complexes containing a cognate A:T base pair as well as A:C and G:T mismatches at the 3’-end (Table 1 and Figure 1D-H).

**Figure 1.**
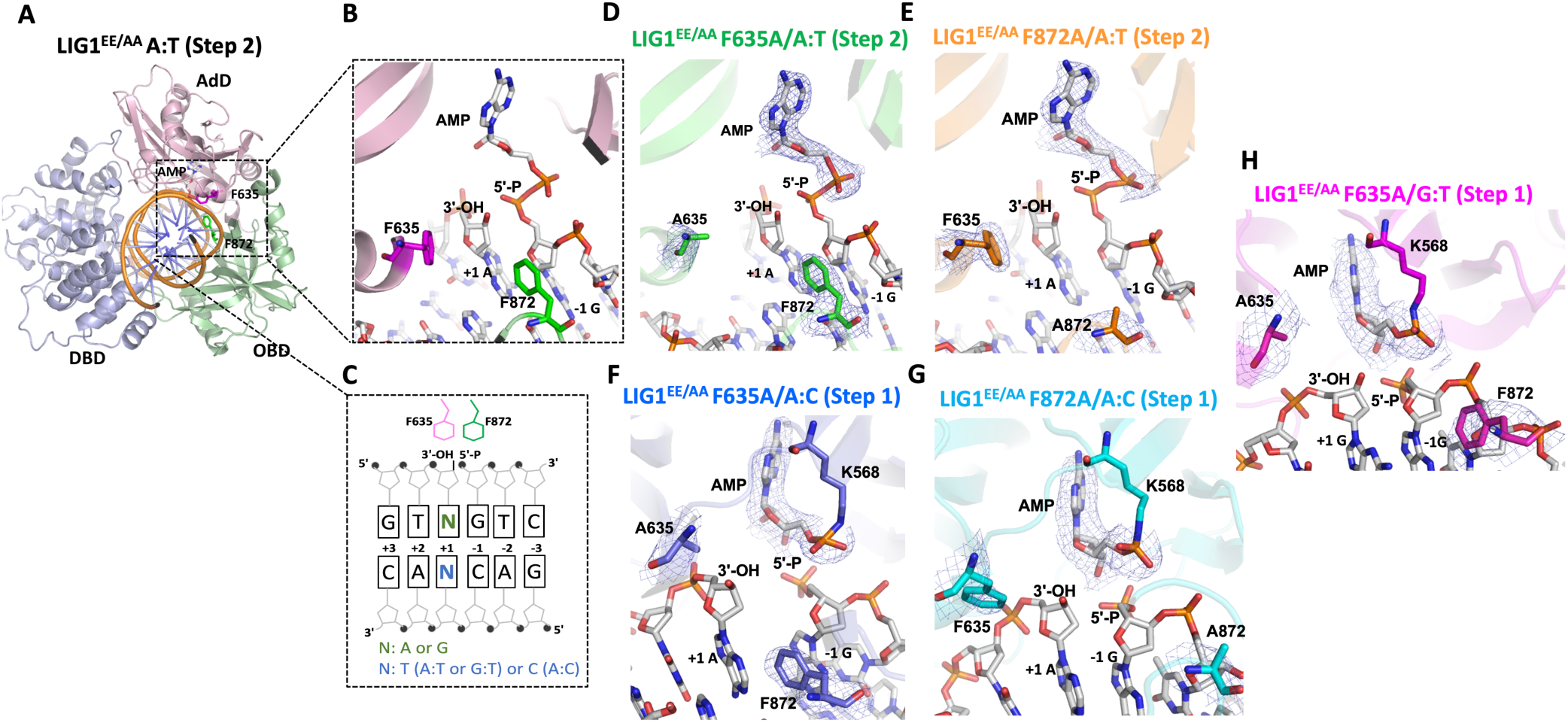
Structures of LIG1^EE/AA^ active site mutants F635A and F872A bound to nick DNA duplexes containing A:T, A:C, and G:T at the 3’-strand. (**A**) Overall structure of LIG1^EE/AA^ (PDB ID: 7SUM) shows that the DBD, AdD, and OBD domains of the protein encircle the nick DNA, in which (**B**) shows the structural details of F635, F872, 3’-OH, and 5’-P at the nick DNA, and (**C**) shows the schematic view of F635, F872, 3’-OH, and 5’-P at the nick DNA. Crystal structure of LIG1 mutants with cognate or mismatched nick DNA duplexes, which are LIG1^EE/AA^ F635A/A:T (step 2, **D**), LIG1^EE/AA^ F872A/A:T (step 2, **E**), LIG1^EE/AA^ F635A/A:C (step 1, **F**), LIG1^EE/AA^ F872A/A:C (step 1, **G**) and LIG1^EE/AA^ F635A/G:T (step 1, **H**). The 2Fo - Fc density map of the AMP, F635/A635, and F872/A872 are contoured at 1.5σ (blue). LIG1 is shown in cartoon mode and the DNA, AMP, F635/A635, and F872/A872 are shown in stick mode.

In the structures of LIG1^EE/AA^ F635A and LIG1^EE/AA^ F872A with A:T nick DNA, we observed that the adenylate moiety has been transferred to the 5’-end of the nick where the DNA-AMP intermediate is formed, which refers to step 2 of ligation reaction (Figure 1D,E). However, in the structures of LIG1^EE/AA^ F635A and LIG1^EE/AA^ F872A with A:C nick DNA, we showed that the ligase can discriminate against this mismatch during step 1 of the ligation reaction where the active site lysine residue (K568) stays adenylated (Figure 1F,G). We previously reported similar differences between A:T and A:C structures of LIG1^EE/AA^ (26). However, in contrast to the mutagenic nick sealing of the G:T mismatch and step 2 structure of LIG1^EE/AA^ as we previously solved (26), the active site mutant structure of LIG1^EE/AA^ F635A bound to a nick DNA duplex containing a G:T mismatch exhibits step 1, suggesting the critical role of F635 for discrimination against incorrect base pairs at the 3’-end (Figure 1H).

### Importance of LIG1 F635 and F872 residues for the discrimination against an oxidatively damaged DNA end

We next solved the structures of F635A and F872A mutants in complex with nick DNA containing 8oxoG:A at the 3’-end (Table 1 and Figure 2). In the structure of LIG1^EE/AA^ F635A/8oxoG:A, we observed that the adenylate has been transferred to the 5’-end of the nick during step 2 of ligation reaction (Figure 2B), which is similar to the previously solved structure of LIG1^EE/AA^ (23). We did not observe any significant differences in the position of the 3’- or 5’-end (Figure 2B). However, the structure of LIG1^EE/AA^ F872A/8oxoG:A was captured during step 1 of the ligation reaction where the active site lysine residue remained adenylated (Figure 2C). Additionally, we observed a shift in the DNA downstream of the nick, which causes a resultant shift away from F635 upstream of the nick (Figure 2C). These results suggest that when the 3’-end is allowed more flexibility, by F635A mutation, the Hoogsteen base pairing of 8oxoG:A could be strong enough to maintain the rigidity that is necessary for adenylate transfer from step 1 to 2 of the ligation reaction. However, when the 5’-end is allowed more flexibility, by F872A mutation, the 8oxoG:A Hoogsteen base pair is not capable of rescuing the conformational shift upstream of the nick caused by the flexibility downstream of the nick.

**Figure 2.**
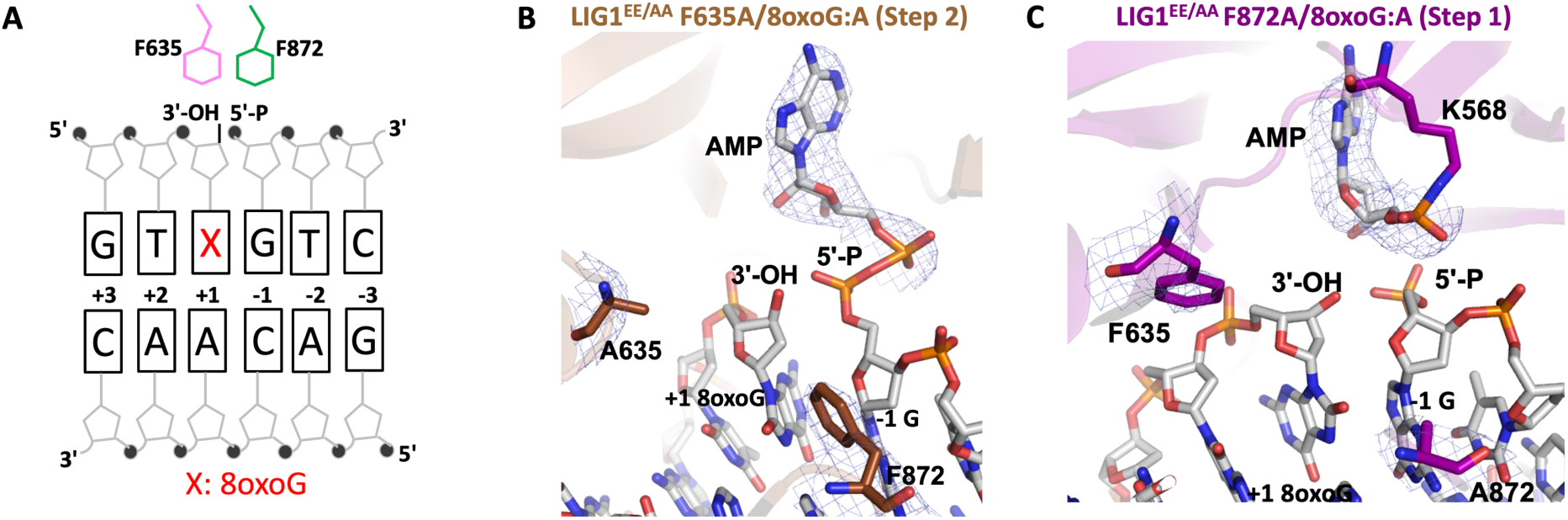
Structures of LIG1^EE/AA^ active site mutants F635A and F872A bound to nick DNA duplex containing 8oxoG:A at the 3’-strand. (**A**) Schematic view of F635, F872, 3’-OH, and 5’-P at the nick DNA. Crystal structure of LIG1^EE/AA^ F635A/8oxoG:A (step 2, **B**) and LIG1^EE/AA^ F872A/8oxoG:A (step 1, **C**). The 2Fo - Fc density map of the AMP, F635/A635, and F872/A872 are contoured at 1.5σ (blue). LIG1 is shown in cartoon mode and the DNA, AMP, F635/A635, and F872/A872 are shown in stick mode.

### The importance of LIG1 active site residues for rigidity of DNA ends at the nick

The overlay of LIG1^EE/AA^ structures for F635A and F872A mutants revealed that alanine substitution at both active site residues creates a small pocket of missing density at the 3’- and 5’-ends, respectively, which allows the ends of the nick more flexibility (Figure 3). This results in displacement of the 3’-end in the structure of LIG1^EE/AA^ F635A/A:C towards the missing density (Figure 3A), increasing the distance between 3’- and 5’-ends as compared to the A:T structures (Figure 3B). In the structure of LIG1^EE/AA^ F635A/G:T, we observed a shift at the 3’-end away from A635, suggesting that the removal of the rigidity imposed by F635 allows distortions in the position of the 3’-end in either direction (Figure 3C). The shift at the 3’-end is accompanied by a shift at the 5’-end of the nick and in the loop containing F872. We did not observe a shift at the 3’-end in the structure of LIG1^EE/AA^ F635A/8oxoG:A or LIG1^EE/AA^ F635A/A:T, suggesting that Hoogsteen or canonical base pairing, respectively, prevents the 3’-end from shifting into the cavity created by F to A substitution at F635 (Figure 3B,D and Supplementary Figure 2).

**Figure 3.**
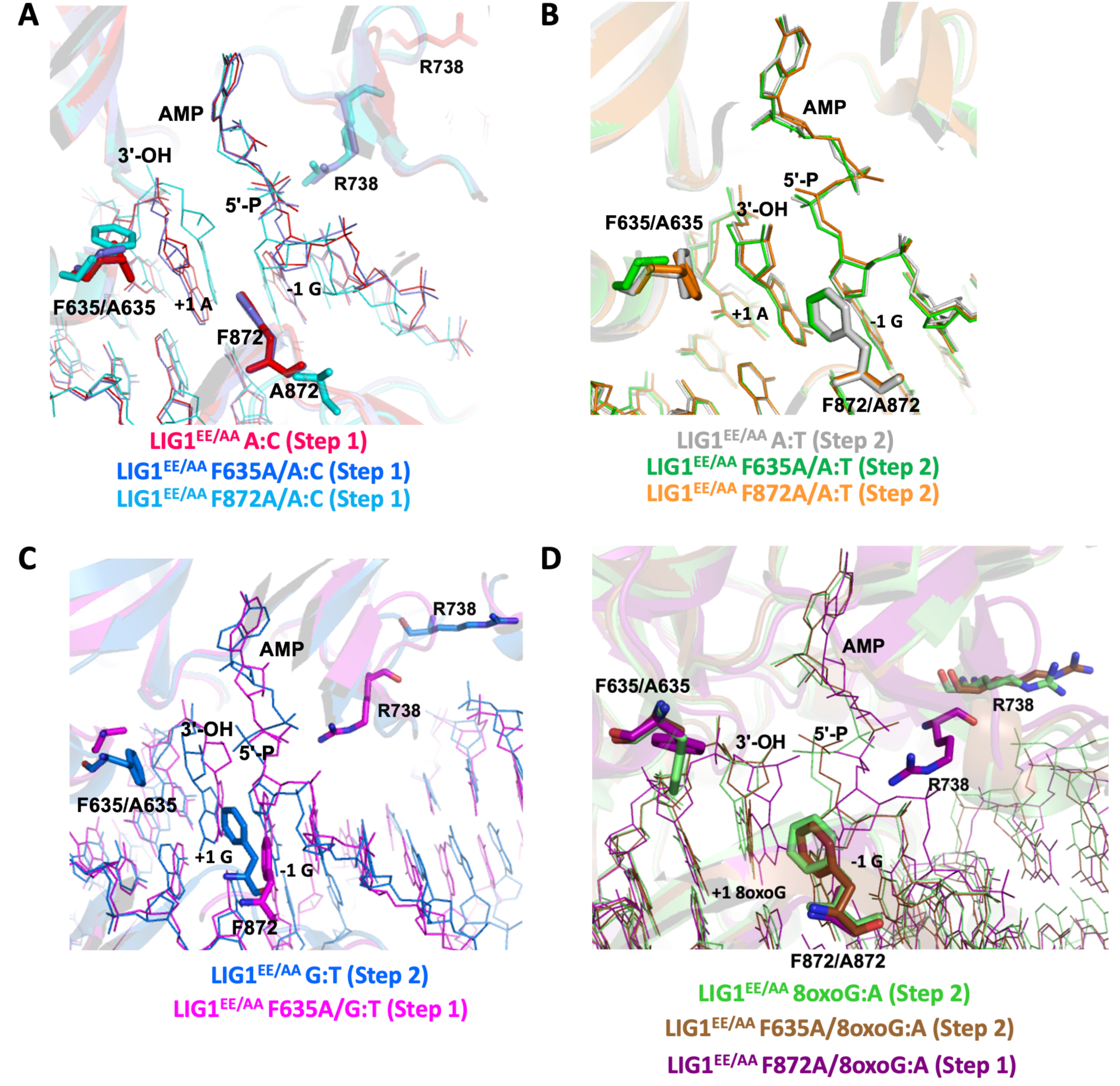
Structures of LIG1^EE/AA^ active site mutants F635A and F872A demonstrate the consequence of loss of DNA end rigidity. (**A**) Overlay of the structures of LIG1^EE/AA^/A:C (PDB ID: 7SX5) with LIG1^EE/AA^ F635A/A:C and LIG1^EE/AA^ F872A/A:C. (**B**) Overlay of the structures of LIG1^EE/AA^/A:T (PDB ID: 7SUM) with LIG1^EE/AA^ F635A/A:T and LIG1^EE/AA^ F872A/A:T. (**C**) Overlay of the structures of LIG1^EE/AA^/G:T (PDB ID: 7SXE) with LIG1^EE/AA^ F635A/G:T. (**D**) Overlay of the structures of LIG1^EE/AA^/8oxoG:A (PDB ID: 6P0E) with LIG1^EE/AA^ F635A/8oxoG:A and LIG1^EE/AA^ F872A/8oxoG:A. LIG1 is shown in cartoon mode. DNA and AMP are shown in line mode. F635/A635 and F872/A872 are shown in stick mode.

When the 5’-end is allowed more flexibility, as we showed in the structure of LIG1^EE/AA^ F872A/A:C, the DNA conformation downstream of the nick changes, which in turn causes a shift upstream of the nick where the DNA shifts away from F635 (Figure 3A). This is accompanied by a shift in the loop containing A872 which was also observed in the structure of LIG1^EE/AA^ F635A/G:T (Figure 3A,C). Similarly, these shifts were not observed in the structure of LIG1^EE/AA^ F872A/A:T, which suggests that canonical base pairing is capable of deterring DNA conformational changes when the 5’-end is allowed more flexibility (Figure 3B). However, unlike our LIG1^EE/AA^ F635A/8oxoG:A structure, in our LIG1^EE/AA^ F872A/8oxoG:A structure, we observed a distortion in the 5’-end of the nick which caused the 3’-end to shift away from F635, suggesting in this instance that Hoogsteen base pairing is not strong enough to enforce the geometric constraints for adenylate transfer (Figure 3D).

The overlay of our current and previously solved structures of LIG1^EE/AA^, grouped into step 1 and step 2 structures, demonstrates the DNA end configuration that enables the ligation reaction to move forward from step 1 to 2 (Supplementary Figure 1) (26). DNA end misalignment, caused by F635A or F872A mutation combined with non-canonical 3’-base pairing, leads to conformational shifts at the DNA ends which prevents this transition between ligation steps. Interestingly, the position of the bound adenosine moiety of the AMP does not show any difference between LIG1 structures that were captured in steps 1 and 2 (Supplementary Figure 1A,D).

### Structures of LIG1 F635A and F872A uncover a shift in the flexible loop that deters mutagenic ligation of A:C and G:T mismatches

We then analyzed the overlay of all our structures near the 5’-end of the nick for both LIG1 mutants (Figure 4A). Interestingly, in all our structures captured in step 1, there was a shift in the flexible loop between residues 733 and 741, especially the guanidinium moiety of R738 which translocates about 15 Å to interact with the 5’-P of the nick (Figure 4B). This shift was not observed in any of our step 2 structures, nor in our previously solved LIG1 structures with cognate A:T or mismatches A:C and G:T as well as all other LIG1 structures reported (Figure 4C) (22–26). Our step 1 structures revealed the putative formation of additional hydrogen bonds between R738 and the 5’-P which would deter AMP transfer from the adenylated lysine residue (K568) to the 5’-end of the nick by increasing the energetic barrier of this transfer (Figure 4B). We suggest that the flexibility of the DNA allows for stabilization of a conformation where the flexible R738 loop interacts with the 5’-P of the nick which would normally not be observed when end rigidity is enforced.

**Figure 4.**
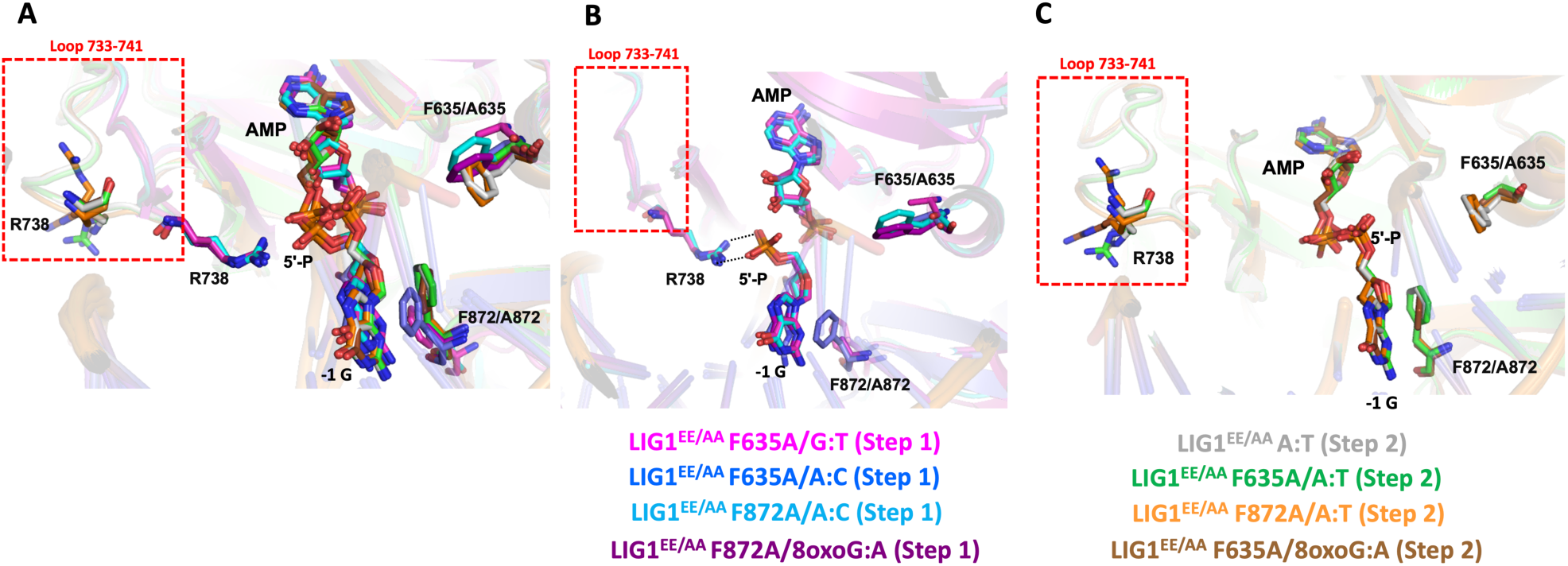
Structures of LIG1^EE/AA^ active site mutants F635A and F872A reveal a flexible loop region. (**A**) Overlay of the structures of LIG1^EE/AA^/A:T (step 2, PDB ID: 7SUM) with LIG1^EE/AA^ F635A/A:T (step 2), LIG1^EE/AA^ F872A/A:T (step 2), LIG1^EE/AA^ F635A/A:C (step 1), LIG1^EE/AA^ F872A/A:C (step 1), LIG1^EE/AA^ F635A/G:T (step 1), LIG1^EE/AA^ F635A/8oxoG:A (step 2), and LIG1^EE/AA^ F872A/8oxoG:A (step 1), in which the loop 733 aa −741 aa has a conformation change shown by a red frame. (**B**) Overlay of the structures of LIG1 mutants with cognate or mismatched nick DNA duplexes in step 2 (LIG1^EE/AA^/A:T, LIG1^EE/AA^ F635A/A:T, LIG1^EE/AA^ F872A/A:T and LIG1^EE/AA^ F635A/8oxoG:A), in which loop 733 aa - 741 aa stays far away from the 3’- and 5’-ends of nick DNA, especially R738 in this loop. (**C**) Overlay of the structures of LIG1 mutants with cognate or mismatched nick DNA duplexes in step 1 (LIG1^EE/AA^ F635A/A:C, LIG1^EE/AA^ F872A/A:C, LIG1^EE/AA^ F635A/G:T and LIG1^EE/AA^ F872A/8oxoG:A), in which the loop 733 aa - 741 aa moves closer to the 5’-end of nick DNA, especially R738, which interacts with the 5’-P of nick DNA.

### Base pairing and sugar pucker analyses of the 3’-ends reveal differences in conformations between LIG1^EE/AA^ F635A and F872A structures

We next investigated the base pairing configuration and sugar pucker conformation of the 3’-end in all of our LIG1^EE/AA^ F635A and F872A structures (Supplementary Figure 2). As expected, the 3’-end adopts a canonical Watson-Crick base pair in the LIG1^EE/AA^/A:T structures (Supplementary Figure 2A,B). Consistent with what we and others previously demonstrated, the A:C and G:T mismatches undergo wobble base pairing in the LIG1 active site while 8oxoG:A forms a Hoogsteen base pair, where the 8oxoG moiety rotates around the N-glycosidic bond to the *syn* conformation (Supplementary Figure 2C-G) (23, 26). Sugar pucker analysis revealed that LIG1^EE/AA^ structures captured in step 1 of the ligation reaction were in the C2’-endo conformation, while all our structures captured in step 2 were in the C3’-endo conformation (Supplementary Figure 2A-G). This demonstrates the importance of the A-form transition upstream of the nick on adenylate transfer.

### Impact of LIG1 active side mutations at F635 and F872 on the substrate specificity of nick DNA containing all 12 non-canonical mismatches

To comprehensively investigate the effect of the active site mutations at F635 and F872 on the mismatch substrate specificity of LIG1, we analyzed the ligation profile of four single mutants F635A, F635L, F872A, F872L as well as the triple-mutants LIG1^EE/AA^ F635A and F872A that we used to solve the structures of LIG1 active site mutants. In addition to these LIG1 mutants, we tested the ligation efficiency of the wild-type and low-fidelity (EE/AA) mutant of LIG1. For this purpose, we used the nick DNA substrates containing all 12 non-canonical mismatches at the 3’-end (Supplementary Scheme 3A).

Consistent with our previous reports (31), in the present study, we showed that wild-type LIG1 can ligate the mismatches with different efficiencies (Supplementary Figures 3 and 4). For example, we observed more than 60% of ligation products for the nick DNA substrates containing T:C, C:T, G:T, C:A, and A:A mismatches. In contrast, the nick DNA containing G:A and A:G mismatches showed the lowest ligation efficiency with less than 10% nick sealing products. We also demonstrated that the EE/AA mutation results in lower ligation fidelity, defined as an increase in the ligation efficiency of non-canonical substrates (Supplementary Figures 5 and 6).

In the presence of the single F635A mutation, the ligation efficiency drastically decreased with less than 10% of ligation products for all 12 non-canonical mismatches (Figure 5 and Supplementary Figure 7). Similarly, our results showed that the LIG1 F635L mutant is not capable of nick sealing in the presence of all 12 non-canonical substrates, except the T:C mismatch which shows a slightly higher amount of ligation products than those of the LIG1 F635A mutant (Figure 6 and Supplementary Figure 8). Regarding the point mutations at F872, we also obtained a complete inability for the ligations of all 12 non-canonical mismatches by both F872A (Figure 7 and Supplementary Figure 9) and F872L (Figure 8 and Supplementary Figure 10) single mutants.

**Figure 5.**
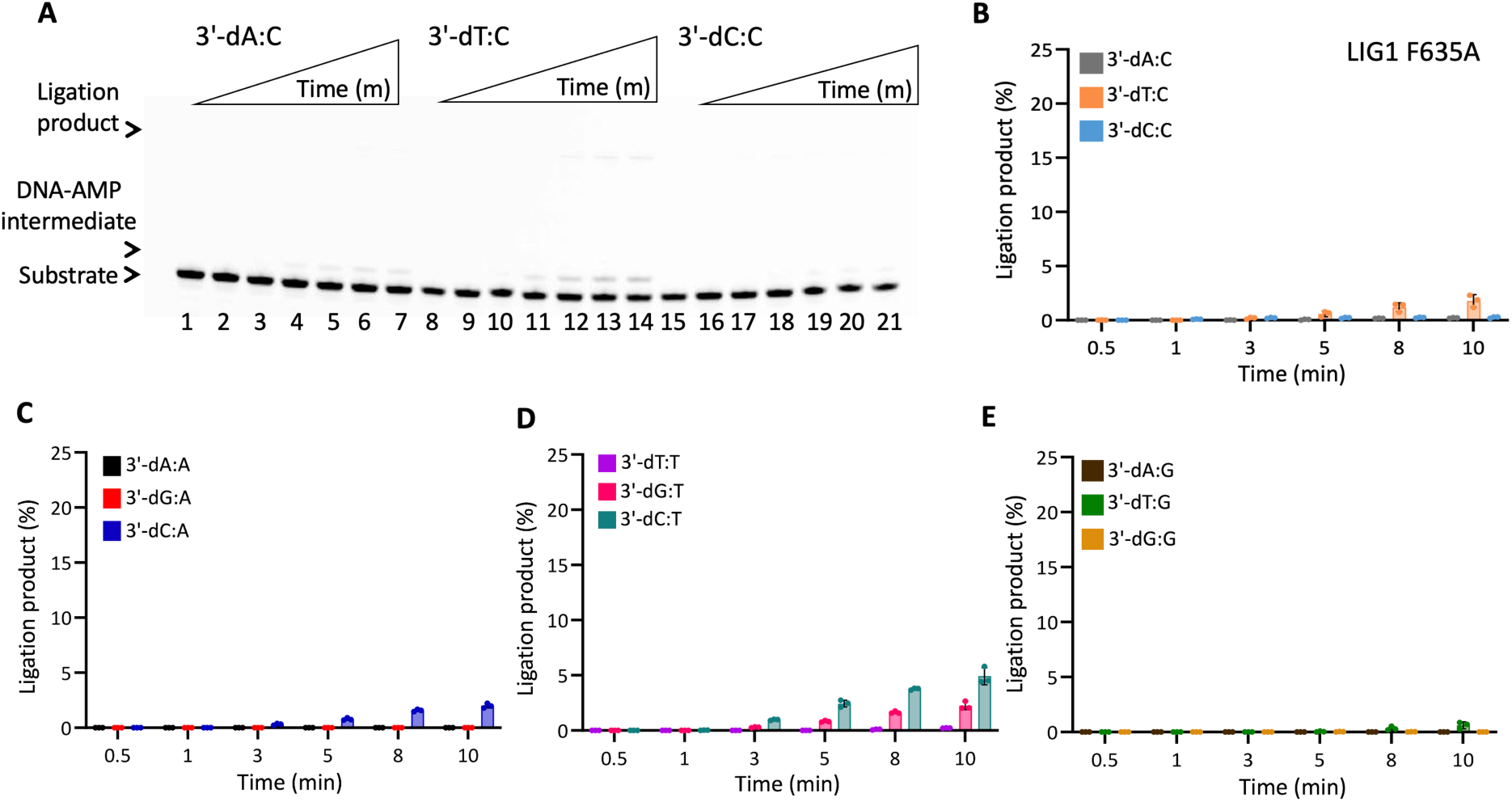
DNA substrate specificity of LIG1 F635A mutant for the nick repair intermediate with all 12 mismatches. (**A**) Lanes 1, 8, and 15 are the negative enzyme controls of the nick DNA substrates containing 3’-preinserted A:C, T:C, and C:C mismatches. Lanes 2-7, 9-14, and 16-21 are the ligation reaction products, and correspond to time points of 0.5, 1, 3, 5, 8, and 10 min. (**B-E**) Graphs show time-dependent changes in the amount of ligation products. The data represent the average from three independent experiments ± SD.

**Figure 6.**
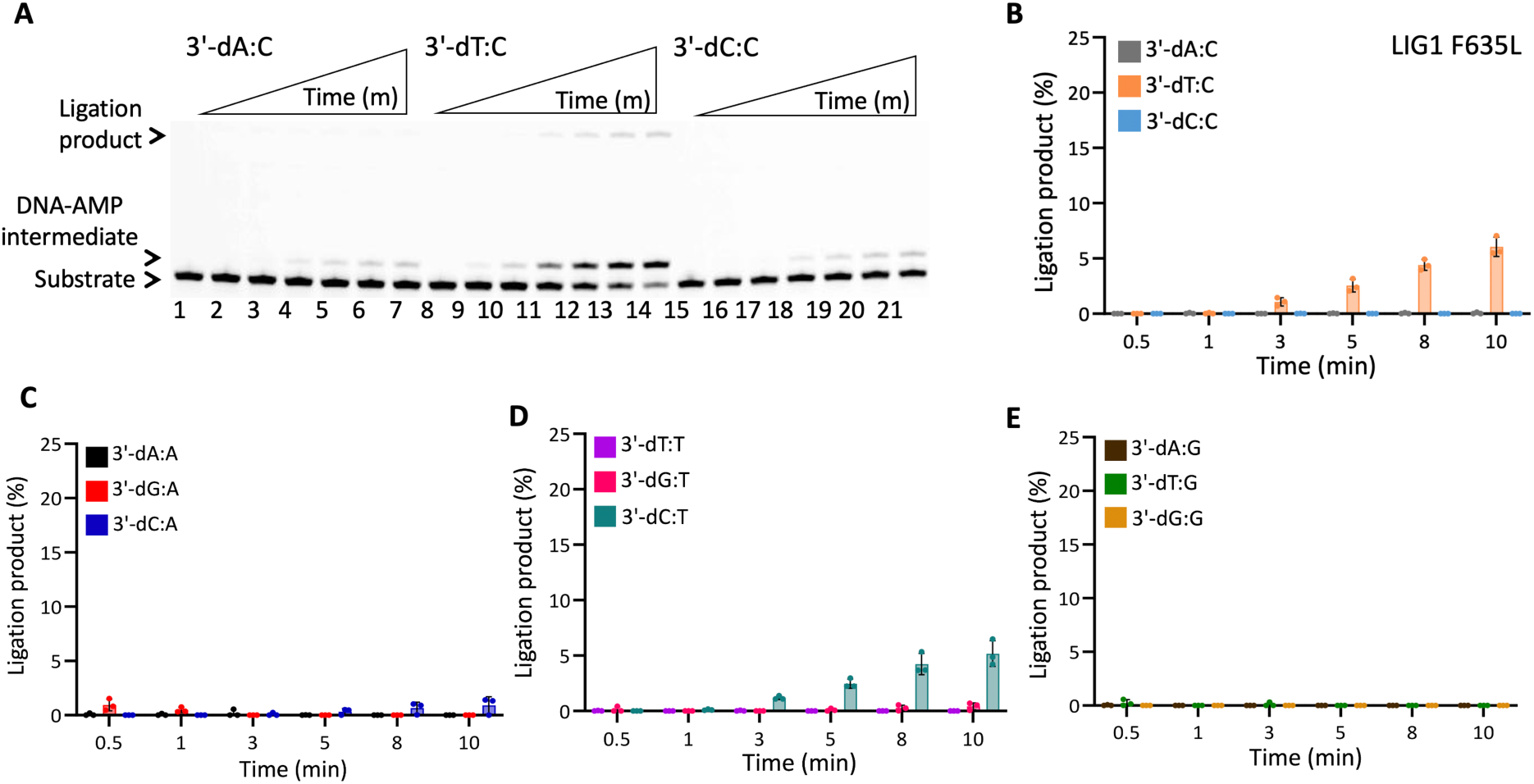
DNA substrate specificity of LIG1 F635L mutant for the nick repair intermediate with all 12 mismatches. (**A**) Lanes 1, 8, and 15 are the negative enzyme controls of the nick DNA substrates containing 3’-preinserted A:C, T:C, and C:C mismatches. Lanes 2-7, 9-14, and 16-21 are the ligation reaction products, and correspond to time points of 0.5, 1, 3, 5, 8, and 10 min. (**B-E**) Graphs show time-dependent changes in the amount of ligation products. The data represent the average from three independent experiments ± SD.

**Figure 7.**
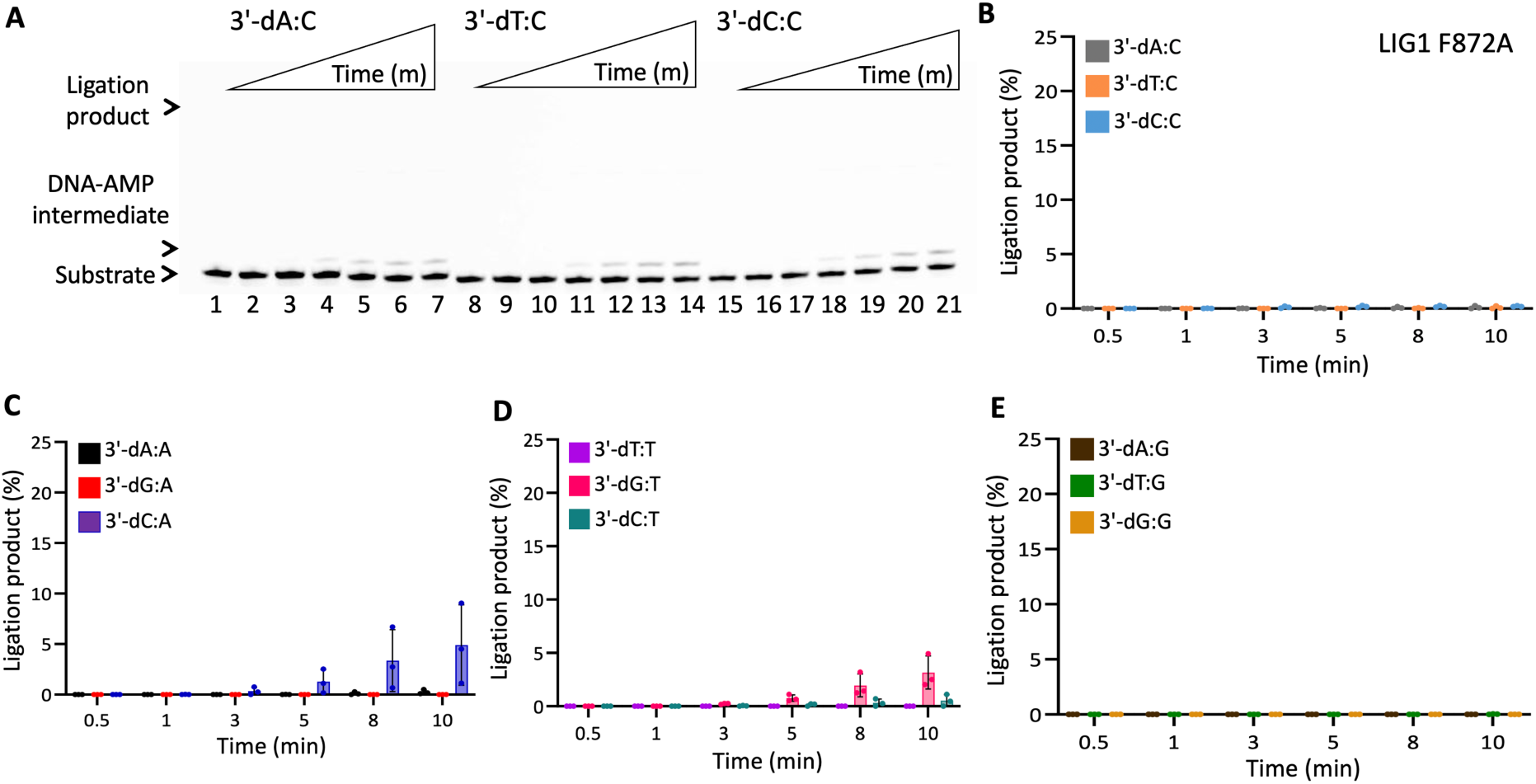
DNA substrate specificity of LIG1 F872A mutant for the nick repair intermediate with all 12 mismatches. (**A**) Lanes 1, 8, and 15 are the negative enzyme controls of the nick DNA substrates containing 3’-preinserted A:C, T:C, and C:C mismatches. Lanes 2-7, 9-14, and 16-21 are the ligation reaction products, and correspond to time points of 0.5, 1, 3, 5, 8, and 10 min. (**B-E**) Graphs show time-dependent changes in the amount of ligation products. The data represent the average from three independent experiments ± SD.

**Figure 8.**
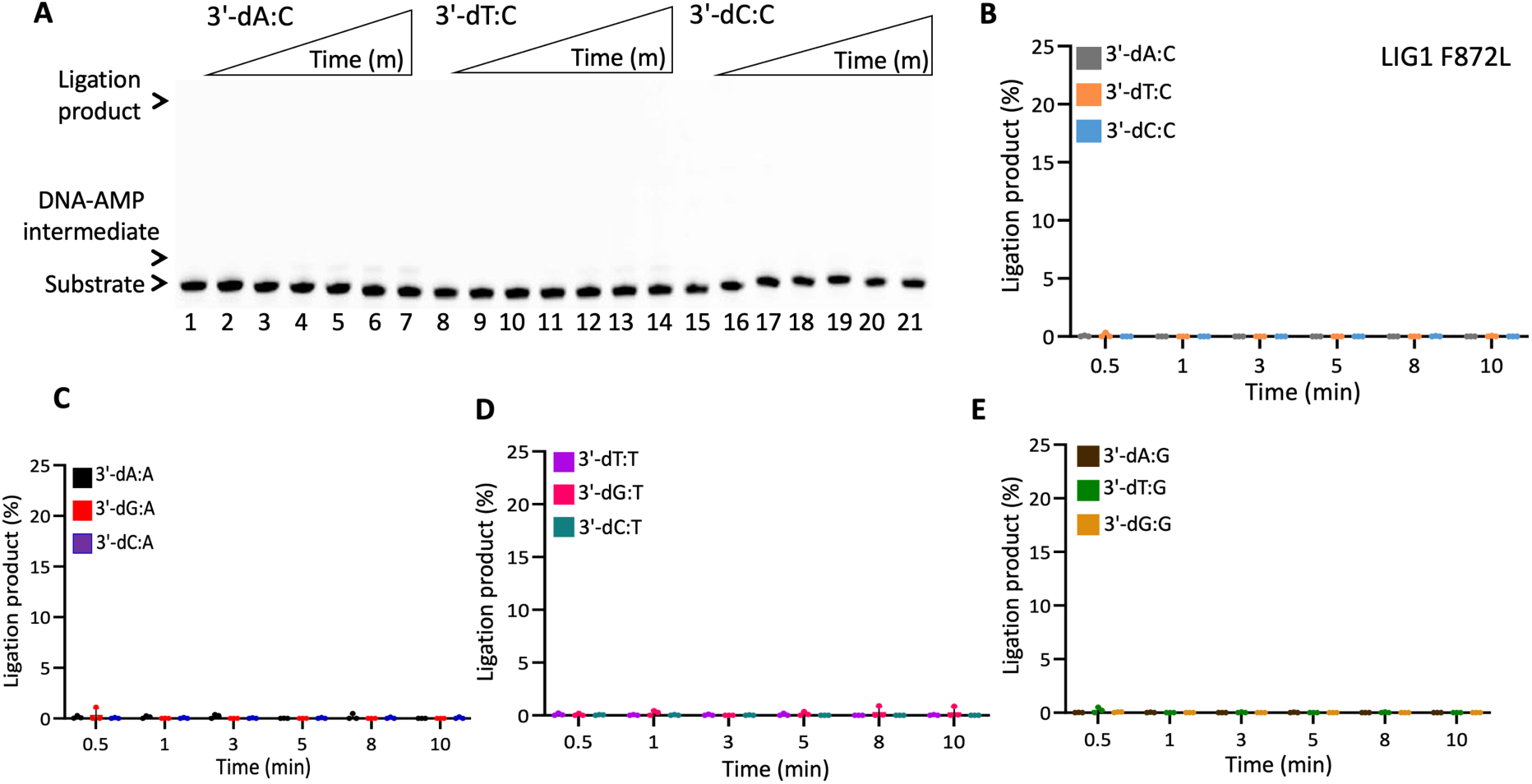
DNA substrate specificity of LIG1 F872L mutant for the nick repair intermediate with all 12 mismatches. (**A**) Lanes 1, 8, and 15 are the negative enzyme controls of the nick DNA substrates containing 3’-preinserted A:C, T:C, and C:C mismatches. Lanes 2-7, 9-14, and 16-21 are the ligation reaction products, and correspond to time points of 0.5, 1, 3, 5, 8, and 10 min. (**B-E**) Graphs show time-dependent changes in the amount of ligation products. The data represent the average from three independent experiments ± SD.

We then questioned whether the EE/AA mutation (LIG1^EE/AA^) that enables LIG1 to seal damaged or mismatched ends more efficiently than the wild-type enzyme can rescue this lack of nick sealing we observed for the single mutants of LIG1 F635A and F872A (23,26,31,33). Interestingly, there was only a very slight rescue in the nick sealing efficiency by the triple mutant of LIG1^EE/AA^ F635A for some of the nick DNA substrates containing 3’-mismatches such as T:C, C:A, and C:T showing 10-20% ligation products (Figure 9 and Supplementary Figure 11). Similarly, we obtained only a minimal rescue between the LIG1^EE/AA^ F872A and the single mutants F872A and F872L for all 12 non-canonical mismatches (Figure 10 and Supplementary Figure 12). For comparison, our results with the LIG1^EE/AA^ mutant showed highly efficient ligation for almost all 12 nick DNA mismatch substrates (Supplementary Figures 5 and 6).

**Figure 9.**
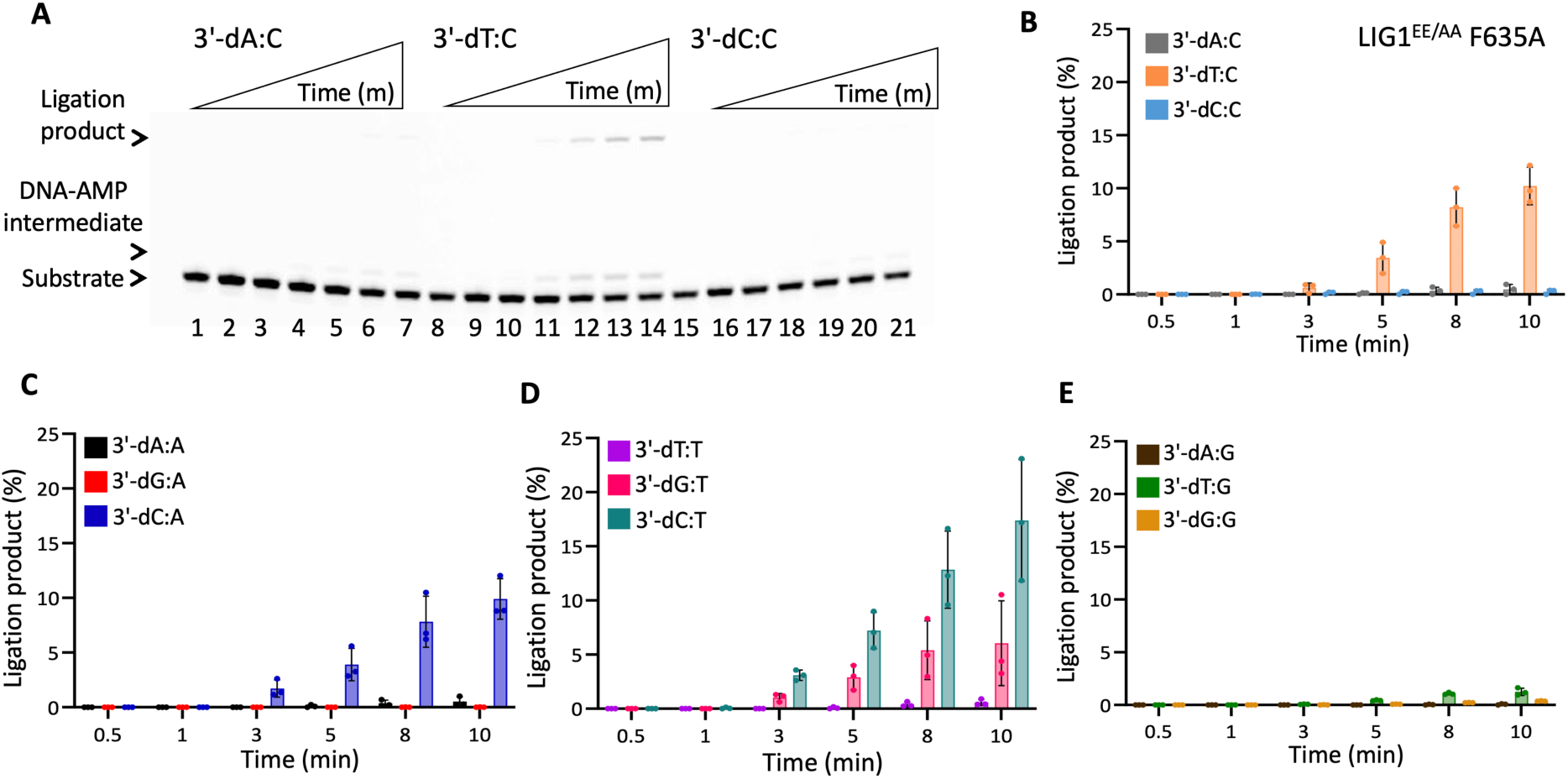
DNA substrate specificity of LIG1^EE/AA^ F635A triple-mutant for the nick repair intermediate with all 12 mismatches. (**A**) Lanes 1, 8, and 15 are the negative enzyme controls of the nick DNA substrates containing 3’-preinserted A:C, T:C, and C:C mismatches. Lanes 2-7, 9-14, and 16-21 are the ligation reaction products, and correspond to time points of 0.5, 1, 3, 5, 8, and 10 min. (**B-E**) Graphs show time-dependent changes in the amount of ligation products. The data represent the average from three independent experiments ± SD.

**Figure 10.**
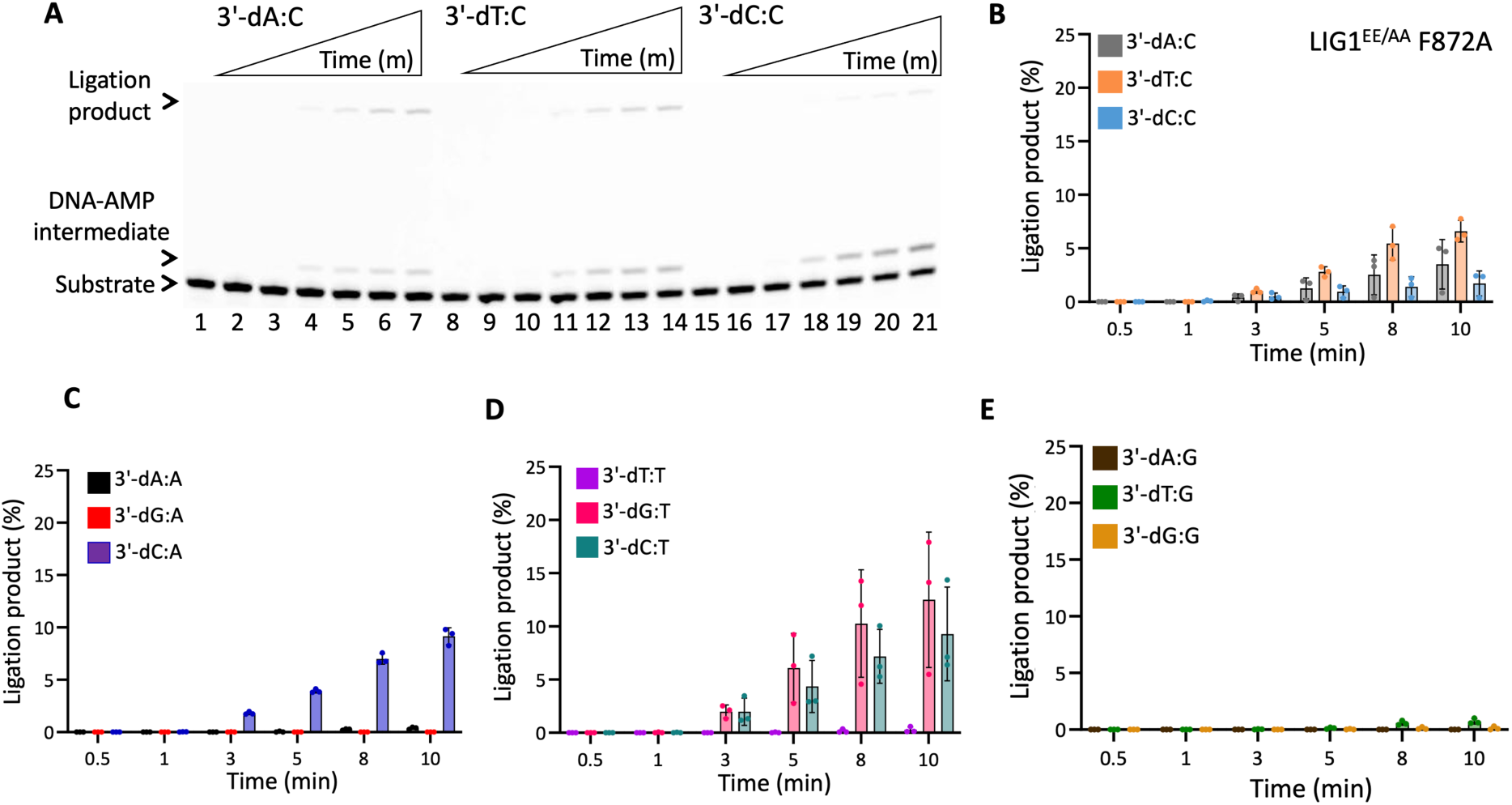
DNA substrate specificity of LIG1^EE/AA^ F872A triple-mutant for the nick repair intermediate with all 12 mismatches. (**A**) Lanes 1, 8, and 15 are the negative enzyme controls of the nick DNA substrates containing 3’-preinserted A:C, T:C, and C:C mismatches. Lanes 2-7, 9-14, and 16-21 are the ligation reaction products, and correspond to time points of 0.5, 1, 3, 5, 8, and 10 min. (**B-E**) Graphs show time-dependent changes in the amount of ligation products. The data represent the average from three independent experiments ± SD.

Finally, we evaluated the nick sealing efficiency of Watson-Crick base paired ends by wild-type LIG1 and all seven LIG1 mutants tested in this study. For this purpose, we used the nick DNA substrates containing 3’-preinserted T:A, A:T, G:C, and C:G. As expected, we obtained robust ligation of these substrates by LIG1 wild-type while there was slightly reduced nick sealing efficiency for F635 and F872 single or triple mutants for all nick DNA substrates containing cognate base pairs (Supplementary Figures 13 and 14). Overall, these results demonstrate that both F635 and F872 residues play critical roles for mismatch discrimination and do not show any substrate specificity such as a type of mismatch or DNA end architecture (Figure 11). In general, we obtained less then 10% ligation for all 12 non-canonical mismatches upon F635 or F872 mutation.

**Figure 11.**
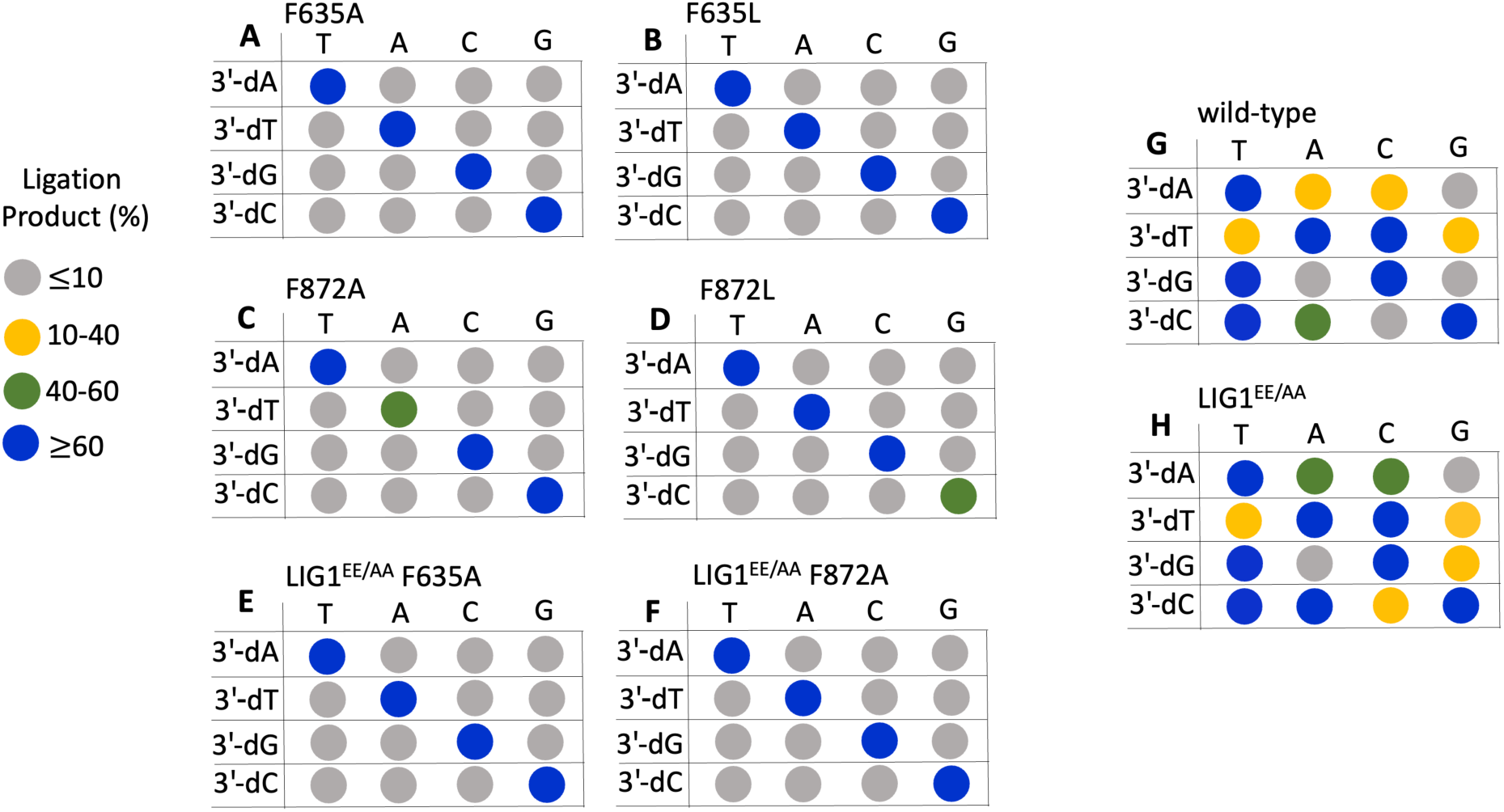
Ligation profile of all 12 non-canonical mismatches by LIG1 wild-type and six active site mutants tested in this study. Dot plots summarizing ligation efficiencies of LIG1 single mutants F635A, F635L, F872A, F872L (**A-D**), triple-mutants LIG1^EE/AA^ F635A (**E**), F872A (**F**), wild-type (**G**), and low fidelity mutant of LIG1^EE/AA^ (**H**) for nick DNA substrates containing all 12 mismatches at the 3’-end. Ligation products (%) represent the data for 3 min.

### Impact of F635 and F872 mutations on the mutagenic ligation of nick DNA with 3’-8oxoG

We next analyzed the ligation efficiency of LIG1 single mutants (F635A, F635L, F872A, F872L) and the triple-mutants LIG1^EE/AA^ F635A and F872A for the nick DNA substrates containing 3’-8oxoG:A and 3’-8oxoG:C to comprehensively understand the effect of the active site mutations on the substrate discrimination against mismatched *versus* damaged DNA ends (Supplementary Scheme 3B).

In line with our previous reports, for the wild-type and LIG1^EE/AA^ mutant of LIG1, we observed the mutagenic ligation of the nick DNA substrate with 3’-8oxoG:A, while the formation of the DNA-AMP intermediate during the ligation reaction is accompanied by mutagenic nick sealing of 3’-8oxoG:C (Supplementary Figures 15 and 16) (30,31). For LIG1 F635A and F635L, nick sealing efficiency is reduced in the presence of 3’-8oxodG:A and we obtained more DNA-AMP intermediates in comparison with wild-type enzyme (Figure 12A-B, lanes 2-7 and Supplementary Figure 16). However, there was greatly diminished ligation with the 3’-8oxoG:C nick substrate (Figure 12A-B, lanes 9-14). For both single mutants, there was time-dependent increase in the amount of ligation products (Figure 12C-D). On the other hand, for the LIG1 mutants F872A and F872L, we did not observe any ligation products for both DNA substrates and relatively less DNA-AMP products accumulated during the ligation reaction (Figure 13 and Supplementary Figure 16). For the triple mutants LIG1^EE/AA^ F635A and F872A, we obtained similar results with those of single mutants (Figure 14). The comparison of ligation products between F635 and F872 mutants *versus* wild-type demonstrates that the low-fidelity mutation of LIG1 (EE/AA) increases the efficiency of 8oxoG:A ligation when F635 is mutated but cannot rescue the lack of ligation for 8oxoG:A when F872A is mutated (Supplementary Figure 17). This is possibly explained by position of the Mg^HiFi^ site which is located upstream of the nick.

**Figure 12.**
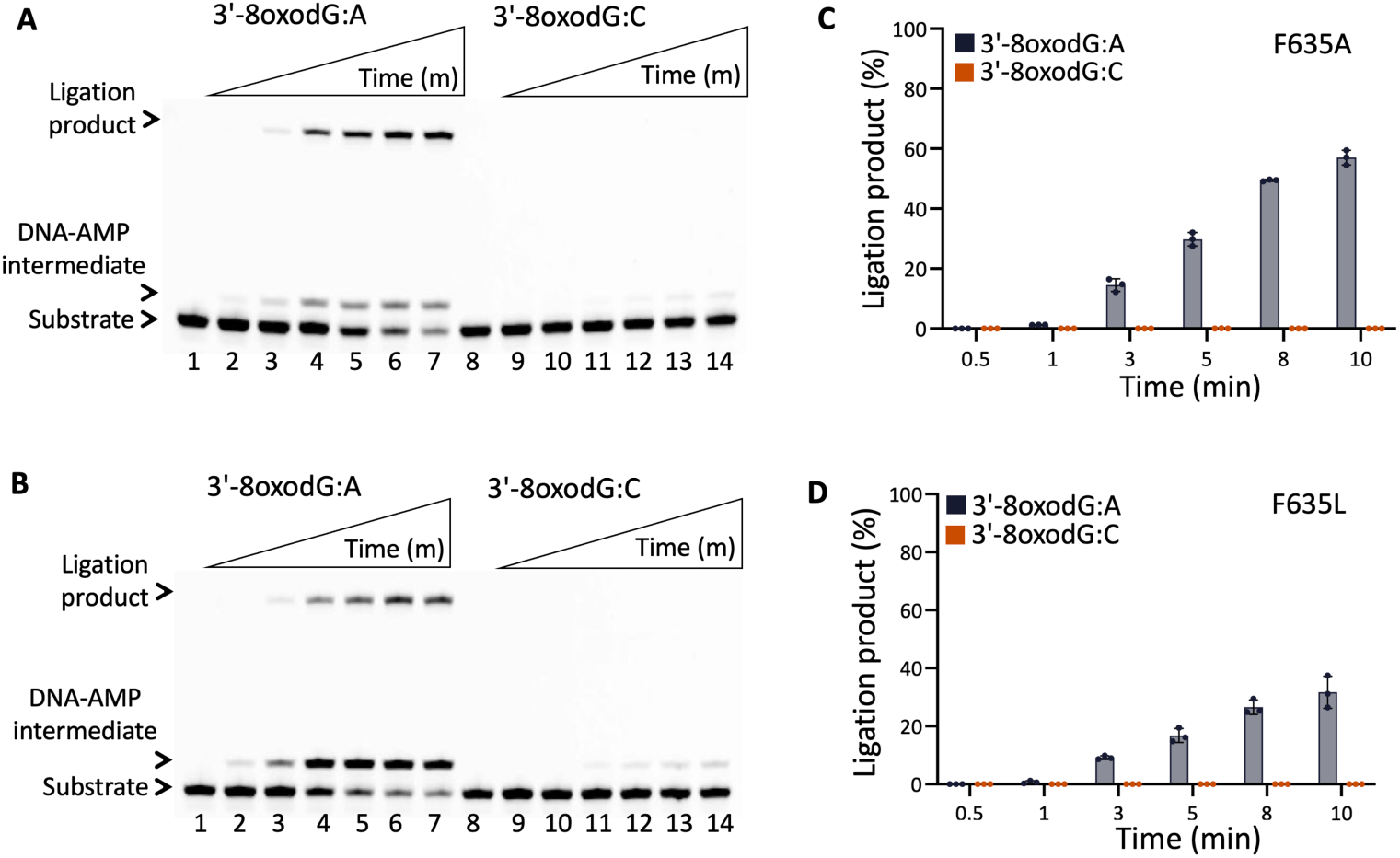
Ligation efficiency of LIG1 F635A and F635L mutants for the nick repair intermediate with 3’-8oxoG. (**A,B**) Lanes 1 and 8 are the negative enzyme controls of the nick DNA substrates containing 3’-8oxodG:A and 3’-8oxodG:C, respectively. Lanes 2-7 and 9-14 are the ligation reaction products by F635A (**A**) and F635L (**B**) single mutants, and correspond to time points of 0.5, 1, 3, 5, 8, and 10 min. (**C,D**) Graphs show time-dependent changes in the amount of ligation products. The data represent the average from three independent experiments ± SD.

**Figure 13.**
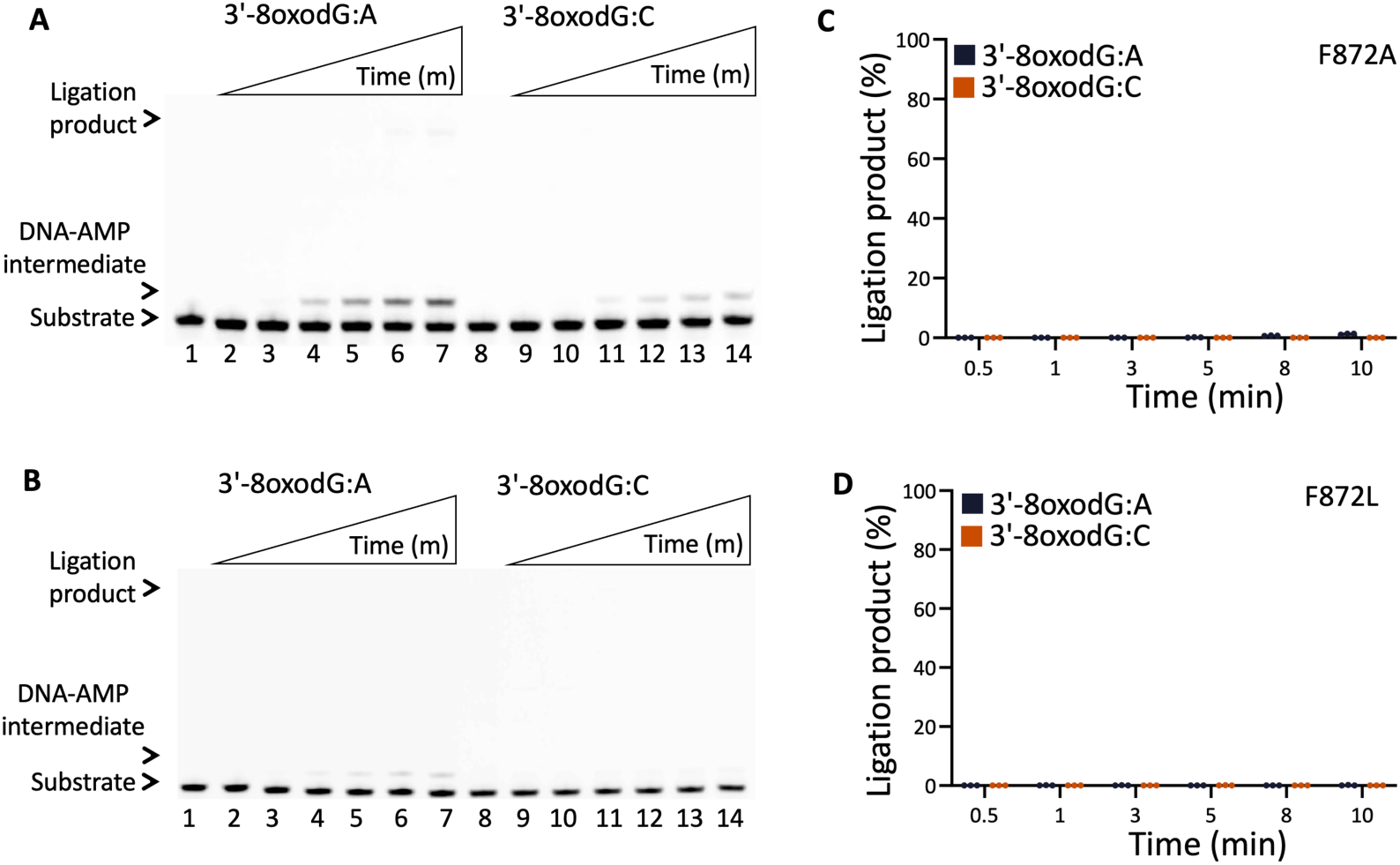
Ligation efficiency of LIG1 F872A and F872L mutants for the nick repair intermediate with 3’-8oxoG. (**A,B**) Lanes 1 and 8 are the negative enzyme controls of the nick DNA substrates containing 3’-8oxodG:A and 3’-8oxodG:C, respectively. Lanes 2-7 and 9-14 are the ligation reaction products by F872A (**A**) and F872L (**B**) single mutants, and correspond to time points of 0.5, 1, 3, 5, 8, and 10 min. (**C,D**) Graphs show time-dependent changes in the amount of ligation products. The data represent the average from three independent experiments ± SD.

**Figure 14.**
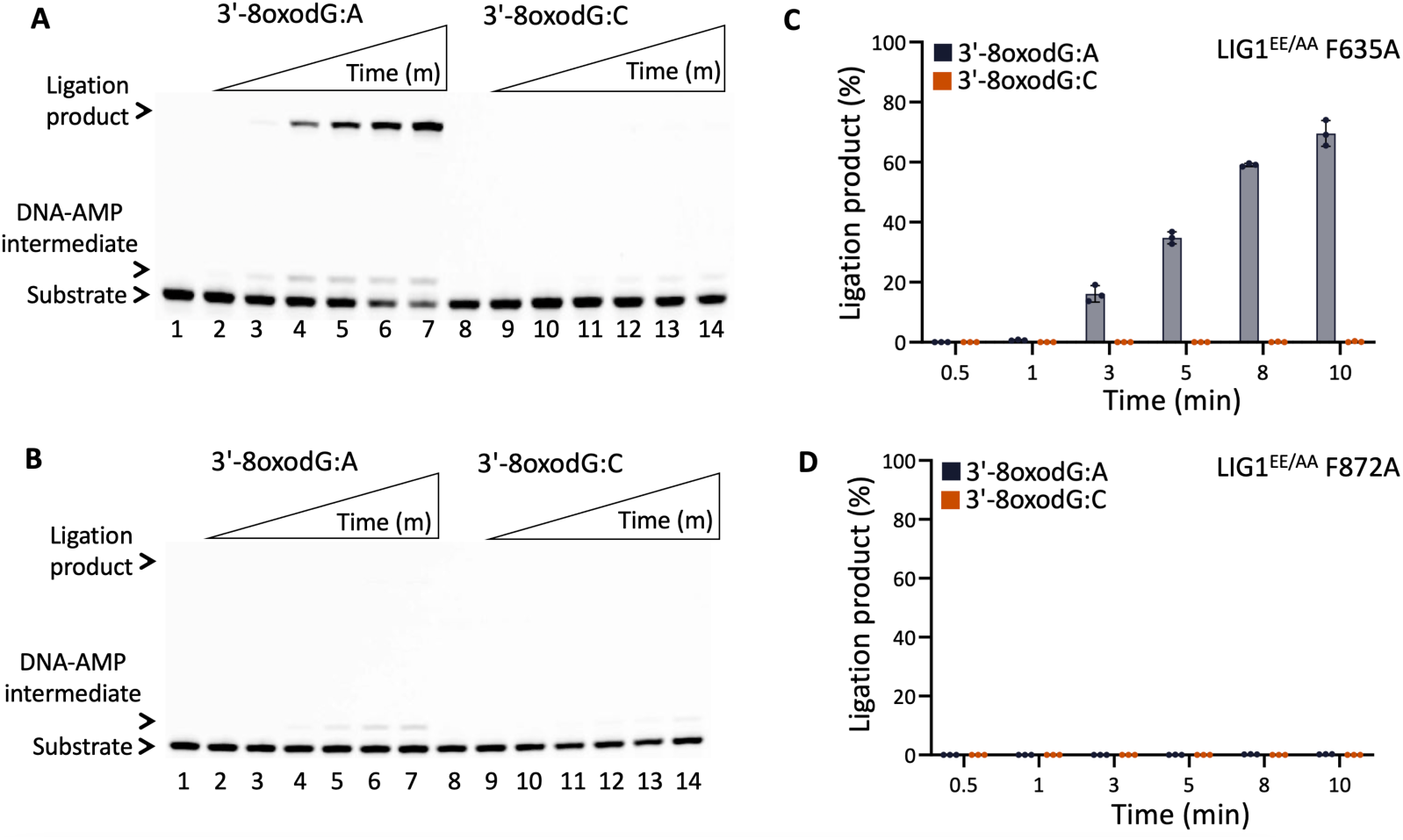
Ligation efficiency of LIG1^EE/AA^ F635A and F872A triple mutants for the nick repair intermediate with 3’-8oxoG. (**A,B**) Lanes 1 and 8 are the negative enzyme controls of the nick DNA substrates containing 3’-8oxodG:A and 3’-8oxodG:C, respectively. Lanes 2-7 and 9-14 are the ligation reaction products by LIG1^EE/AA^ F635A (**A**) and F872A (**B**) triple mutants, and correspond to time points of 0.5, 1, 3, 5, 8, and 10 min. (**C,D**) Graphs show time-dependent changes in the amount of ligation products. The data represent the average from three independent experiments ± SD.

### Effect of the active site mutations on LIG1 stability and folding

We finally evaluated whether the LIG1 mutations at F635 and F872 impact protein stability and folding by thermal shift protein denaturation assays. For this purpose, we performed the assays in the presence and absence of nick DNA containing 3’-preinserted cognate G:C.

Our results demonstrated that all LIG1 proteins tested in this study showed similar T_m_ values, which demonstrates similar thermal stability indicating similar global protein folds (Supplementary Figure 18A,B). LIG1 F872A and F872L mutants exhibit slightly decreased thermal stability, but not drastic enough to demonstrate misfolding or large disturbances in the protein structure. We also tested the thermal stability of the LIG1 wild-type and mutants in the presence of nick DNA. Our results demonstrated an increase in T_m_ values for all LIG1 variants indicating that all LIG1 variants are able to bind nick DNA and suggesting that DNA binding stabilizes the ligase proteins (Supplementary Figure 18C). We also observed higher thermal stability of both triple mutants suggesting that mutation at the Mg^HiFi^ site leads to higher thermal stability upon nick DNA binding (Supplementary Figure 18).

## Discussion

ATP-dependent DNA ligases catalyze the formation of a phosphodiester bond in a three-step reaction which is accompanied by large-scale rearrangements of the AdD and OBD domains of the ligase to expose the catalytic site for cofactor (ATP or NAD^+^) and nick DNA substrate binding (1–6). Previously solved crystal structures of ATP or NAD^+^-dependent DNA ligases from other sources such as *Chlorella virus*, *Mycobacterium tuberculosis*, *Pyrococcus furiosus*, and *Thermus thermophilus* reported that both base pair geometry of the 3’-base pair and the recognition of minor groove contacts by hydrogen bond interactions between the ligase and the DNA duplex ensures ligation fidelity (38–56). Several reports have described efficient joining at nicks with mismatches by eukaryotic DNA ligases (41–44). For human LIG1 and LIG3, it has been shown that the ligation of 3’-mismatches is delayed depending on ligation conditions such as salt concentration, suggesting that suppression of nick sealing at mismatches could be a cellular surveillance mechanism for reducing spontaneous mutation rates (38). We previously reported that the ligation fidelity at the final step of base excision repair (BER) pathway is sensitive to the DNA-end configuration of nick repair intermediates and that the identity of 3’-mismatch compromises the efficiency of nick sealing distinctly by BER ligases I and IIIα/XRCC1 complex (31).

LIG1 is recognized to be the primary replicative ligase and ligates single-strand DNA breaks during DNA repair pathways (12–20). X-ray structures of LIG1 demonstrated a unique Mg^2+^-mediated fidelity mechanism by which the rigidity imposed by metal binding allows for mismatch discrimination at the 3’-end of the nick (23). Furthermore, it has been proposed that LIG1 mutations associated with the immunodeficiency disease destabilize DNA-ligase interaction by compromising this Mg^2+^ binding affinity (24). In our previously reported LIG1/mismatch structures, we showed that the ligase can distinguish Watson-Crick geometry A:T from mismatches A:C, while the G:T evades discrimination well by engaging the ligase active site in the wobble conformation (26). However, how LIG1 interacts with the DNA duplex through the nature of active site contacts to proofread unusual ends during nick sealing of nuclear replication and DNA repair at atomic resolution is still unknown.

In the present study, we further interrogated the DNA substrate-ligase interaction in the presence of a mismatch or oxidative damage at the 3’-end of nick and characterized the roles of LIG1 conserved active site residues for maintaining ligase fidelity by comprehensive biochemical characterization and X-ray structures. Upon mutation of F635 and F872 to either alanine (A) or leucine (L), we obtained a drastic ablation of nick sealing efficiency for all possible 12 non-canonical mismatches with only a minor loss of catalytic efficiency for canonical DNA ends. Our findings suggest that these active site residues are both required for efficient non-canonical catalysis and that all mutations at these residues (F635A, F635L, F872A, F872L) result in higher fidelity ligation by deterring nick sealing of mutagenic non-canonical ends.

Structures of LIG1^EE/AA^ F635A and F872A mutants in complex with nick DNA duplexes containing a canonical A:T base pair, A:C and G:T mismatches, and 8oxoG:A at the 3’-end revealed further insights into the importance of these active sites for the mismatch discrimination mechanism of LIG1. We observed that the substitution of F to A at both F635 and F872 residues creates a pocket of missing density that the aromatic moiety of phenylalanine would normally occupy, this in turn allows for a greater degree of flexibility at the 3’- and 5’-ends, respectively. This flexibility results in increased sensitivity to the identity of the base-pair at the 3’-terminus which is dependent on the position of a specific point mutation.

When the 3’-end is allowed greater flexibility as a result of F635A mutation, the presence of this missing density, in combination with the improperly base paired mismatch, enables the 3’-end to shift into the pocket, as we observed in the structure of LIG1^EE/AA^ F635A/A:C mismatch. Alternatively, it can shift away from the 3’-end, as we observed in the structure of LIG1^EE/AA^ F635A/G:T, which deters mutagenic nick sealing of the G:T mismatch by causing DNA end misalignment. Our results suggests that the direction of the 3’-end shift is likely mismatch dependent. However, this shift was not observed in the structures of LIG1^EE/AA^ F635A for a cognate A:T and damaged 8oxoG:A ends, which suggests that canonical or Hoogsteen base pairing is strong enough to enforce the active site geometry required for ligation even in the presence of missing sidechain density. When the 5’-end is allowed greater flexibility as a result of F872A mutation, the DNA conformation downstream of the nick changes, resulting in a subsequent change upstream of the nick, and therefore misaligns the DNA ends which provide a barrier for catalysis. Unlike F635A mutation, upon F872A mutation, Hoogsteen base pairing is not capable of enforcing the active site geometry required for efficient catalysis, as we observed in the structure of LIG1^EE/AA^ F872A/8oxoG:A. Only a proper canonical base pair is capable of enforcing the active site geometry required for efficient catalysis, as we observed in the structure of LIG1^EE/AA^ F872A/A:T. These results suggest that the 5’-end stability could be more critical for efficient ligation of non-canonical substrates. Our findings suggest that the conserved LIG1 active site phenylalanine residues, F635 and F872, are critical for nick sealing of non-canonical 3’-ends by enforcing rigidity of the 3’- and 5’-ends, respectively, through interactions with the sugar moiety.

Furthermore, our LIG1 structures that were captured in step 1 of the ligation reaction revealed a shift in a flexible loop region which results in the positioning of the guanidinium moiety of R738 within hydrogen bonding distance of the 5’-P of the nick. These putative hydrogen bonds provide a barrier to adenylate transfer which, in combination with DNA end misalignment, explains our *in vitro* observation of inefficient non-canonical ligation by all single and triple mutations at F635 and F872. It is not immediately obvious why this shift of the flexible loop only manifests upon F635 or F872 mutation, but it is consistent with previous studies which have also shown that mutations at one side of the nick can influence the ligase active site on the other side of the nick and highlights the sensitivity of the active site of LIG1 to mutation (24). Future biochemical and structural studies are required to understand how this shift only manifests upon F635A or F872A mutation and to elucidate the impact of R738 on the mismatch discrimination mechanism of LIG1. Because these phenylalanine residues are well conserved among human and other non-human DNA ligases (39–56), the high-fidelity ligation we observed by the LIG1 mutations at F635 and F872 could be a universal mechanism of 3’-end discrimination that other DNA ligases could employ to maintain fidelity during the ligation step of DNA replication, repair, and recombination.

## Funding

This work was supported by a grant 1R35GM147111-01 from the National Institute of General Medical Sciences (NIGMS).

## Acknowledgements

This research used resources of the Advanced Photon Source (APS), a U.S. Department of Energy (DOE) Office of Science user facility operated for the DOE Office of Science by Argonne National Laboratory under Contract No. DE-AC02-06CH11357. The authors thank Dr. Craig Vander Kooi (University of Florida) for his exceptional support with the APS beamtime and data collection. The authors thank Abigail Ortiz and Ernesto Martinez for help with LIG1 construct preparation and protein purification.

## Data availability

Atomic coordinates and structure factors for the reported crystal structures of LIG1 in complex with nick DNA have been deposited in the RCSB Protein Data Bank under accession number LIG1^EE/AA^ F635A A:T (8EZY), A:C (8F0C), G:T (8GKE), 8oxoG:A (8GKI) and LIG1^EE/AA^ F872A A:T (8GIK), A:C (8GIQ), 8oxoG:A (8GJO). All relevant data are available from the authors upon reasonable request.

## Author contributions

Conceptualization M.Ç., methodology and investigation M.G., T.Q., M.P.; writing-original draft, M.Ç., M.G.; T.Q.; writing-reviewing, editing original draft and revision, M.Ç., M.G.; T.Q.; funding acquisition M.Ç.

## Competing interests

The authors declare no competing interests.

## Additional information

Correspondence and requests for materials should be addressed to Melike Çağlayan.

## Supplementary Information

**Supplementary Figure 1.**
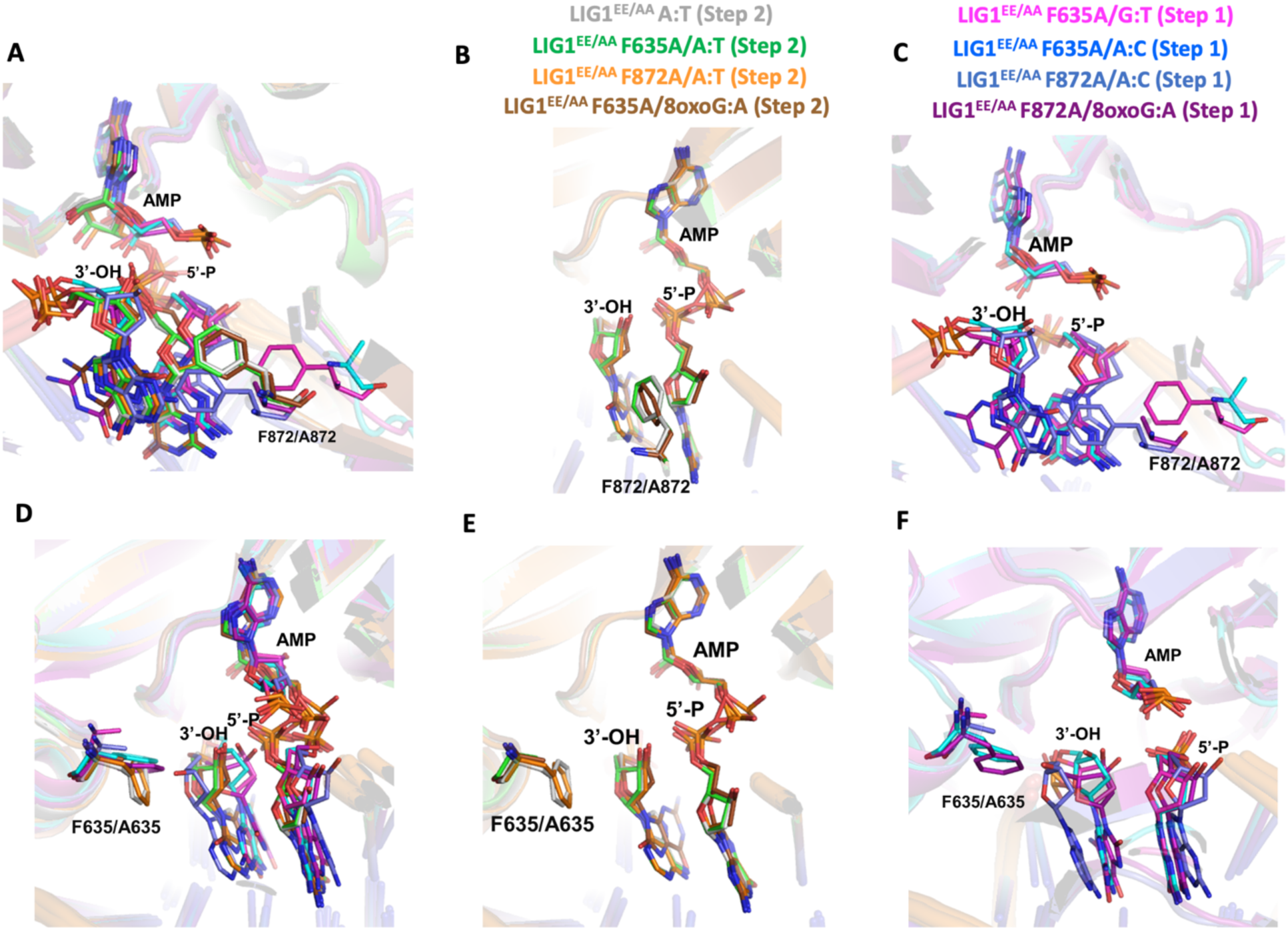
Structures of LIG1^EE/AA^ active site mutants F635A and F872A demonstrate the geometry required for the conversion from step 1 to 2 of the ligation reaction. Overlay of the structures of LIG1^EE/AA^/A:T (PDB ID: 7SUM), LIG1^EE/AA^ F635A/A:T, LIG1^EE/AA^ F635A/A:C, LIG1^EE/AA^ F635A/G:T, LIG1^EE/AA^ F635A/8oxoG:A, LIG1^EE/AA^ F872A/A:T, LIG1^EE/AA^ F872A/A:C, and LIG1^EE/AA^ F872A/8oxoG:A emphasizes the alignment of the DNA ends as well as the positions of F872/A872 (**A**) and F635/A635 (**D**). Overlay of our current and previously solved structures captured in step 2 of the ligation reaction emphasizes the alignment of the DNA ends as well as the positions of F872/A872 (**B**) and F635/A635 (**E**). Overlay of our current structures captured in step 1 of the ligation reaction emphasizes the alignment of the DNA ends as well as the positions of F872/A872 (**C**) and F635/A635 (F).

**Supplementary Figure 2.**
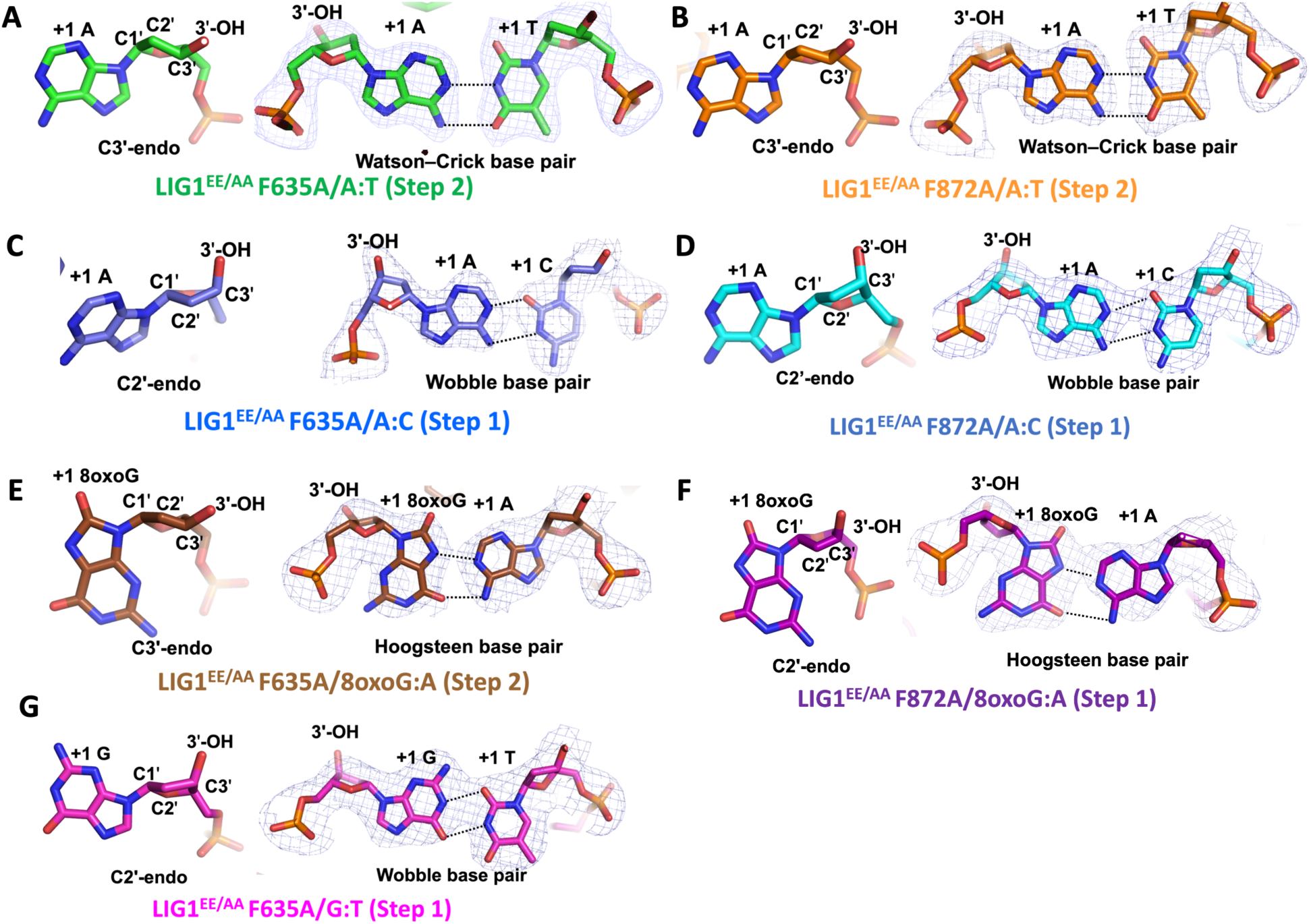
Analysis of the 3’-ends reveals the base pairing and sugar pucker differences between structures of LIG1^EE/AA^ active site mutants F635A and F872A. The sugar pucker and base pairing conformation is shown for LIG1^EE/AA^ F635A/A:T (**A**), LIG1^EE/AA^ F872A/A:T (**B**), LIG1^EE/AA^ F635A/A:C (**C**), LIG1^EE/AA^ F872A/A:C (**D**), LIG1^EE/AA^ F635A/8oxoG:A (**E**), LIG1^EE/AA^ F872A/8oxoG:A (**F**), and LIG1^EE/AA^ F635A/G:T (**G**). The 2Fo - Fc map density map of the 3’-base pair is contoured at 1.5σ (blue).

**Supplementary Figure 3.**
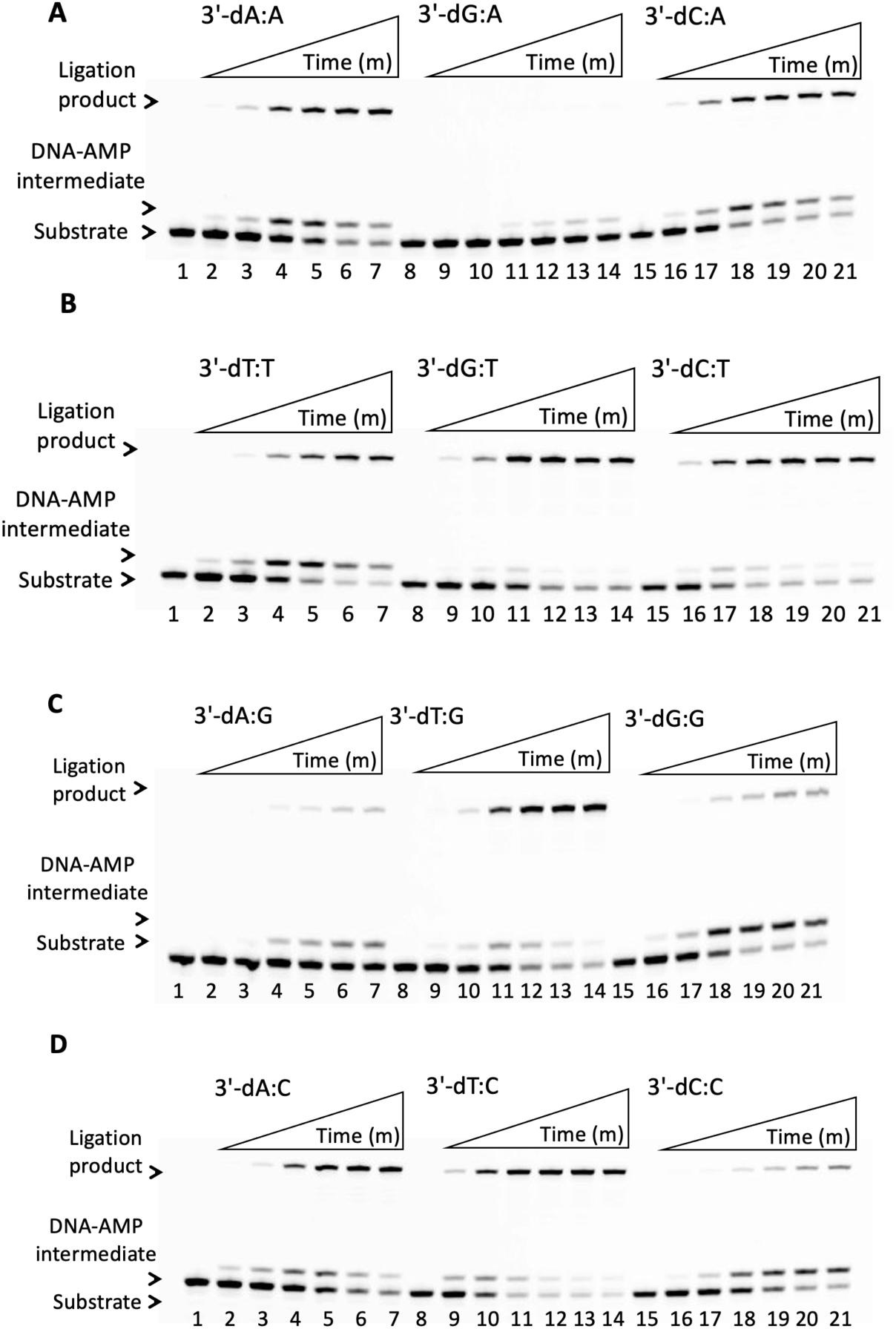
Ligation of nick DNA substrates containing template A, T, G, and C mismatches by wild-type LIG1. (**A-D**) Lanes 1, 8, and 15 are the negative enzyme controls of the nick DNA substrates containing template A, T, G, and C mismatches. Lanes 2-7, 9-14, and 16-21 are the ligation products in the presence of nick DNA substrates with 3’-mismatches by wild-type LIG1, and correspond to time points of 0.5, 1, 3, 5, 8, and 10 min.

**Supplementary Figure 4.**
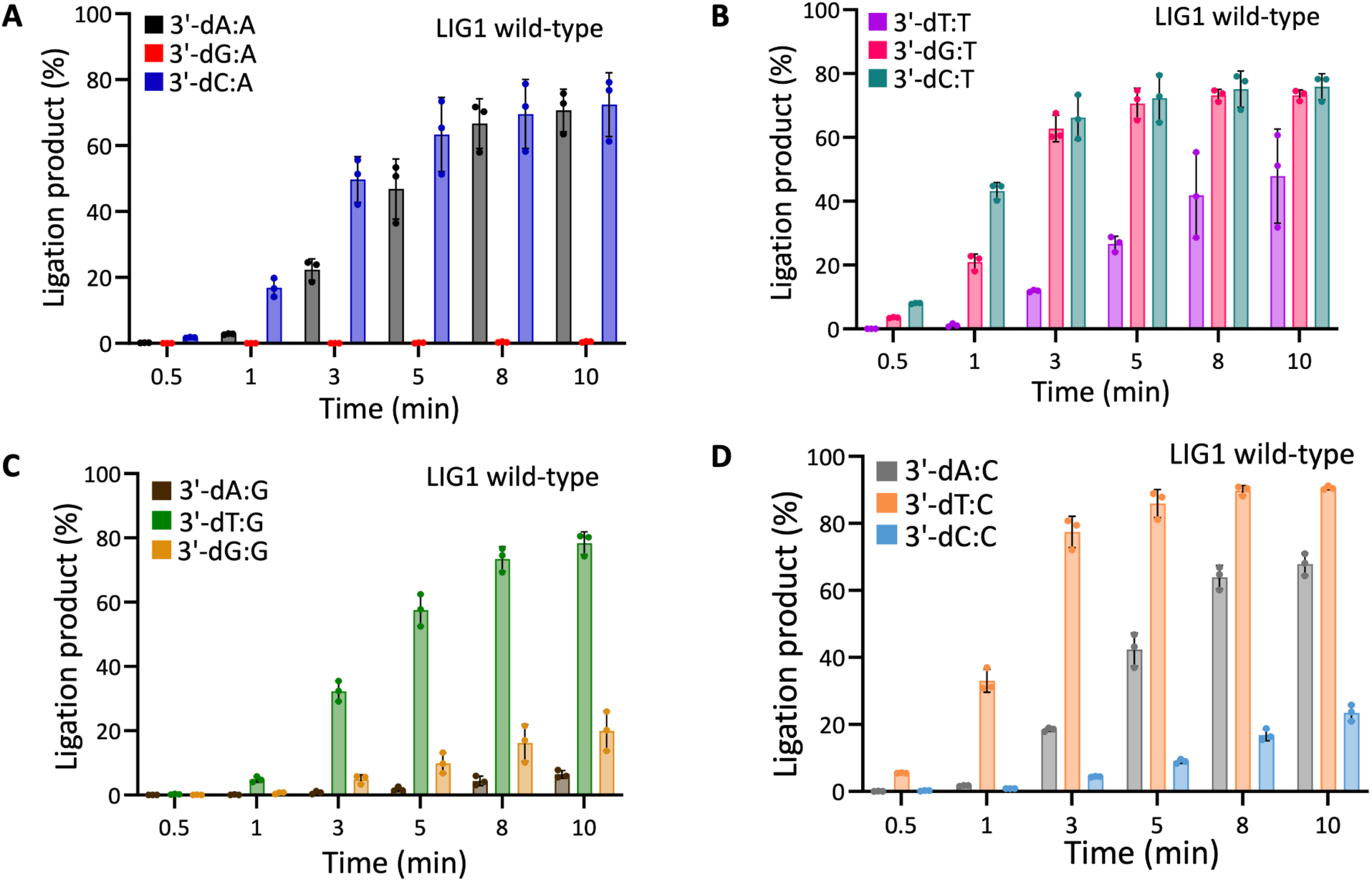
Ligation of nick DNA substrates containing template A, T, G and C mismatches by wild-type LIG1. (**A-D**) Graphs show time-dependent changes in the amounts of ligation products in the presence of template A (A), T (B), G (C), and C (D) mismatches by wild-type LIG1. The data represent the average from three independent experiments ± SD.

**Supplementary Figure 5.**
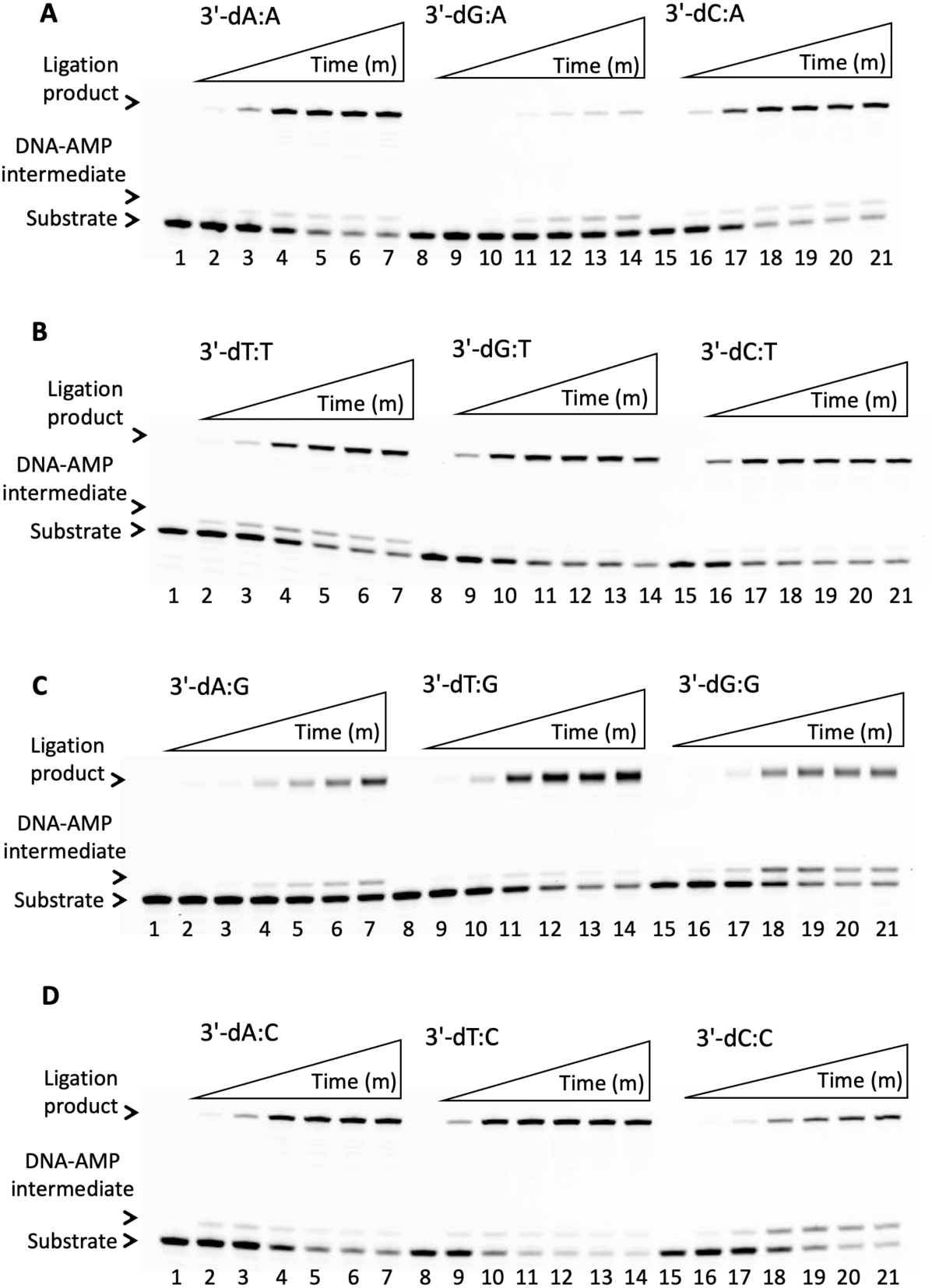
Ligation of nick DNA substrates containing template A, T, G, and C mismatches by LIG1^EE/AA^ double mutant. (**A-D**) Lanes 1, 8, and 15 are the negative enzyme controls of the nick DNA substrates containing template A, T, G, and C mismatches. Lanes 2-7, 9-14, and 16-21 are the ligation products in the presence of nick DNA substrates with 3’-mismatches by LIG1^EE/AA^ low-fidelity mutant, and correspond to time points of 0.5, 1, 3, 5, 8, and 10 min.

**Supplementary Figure 6.**
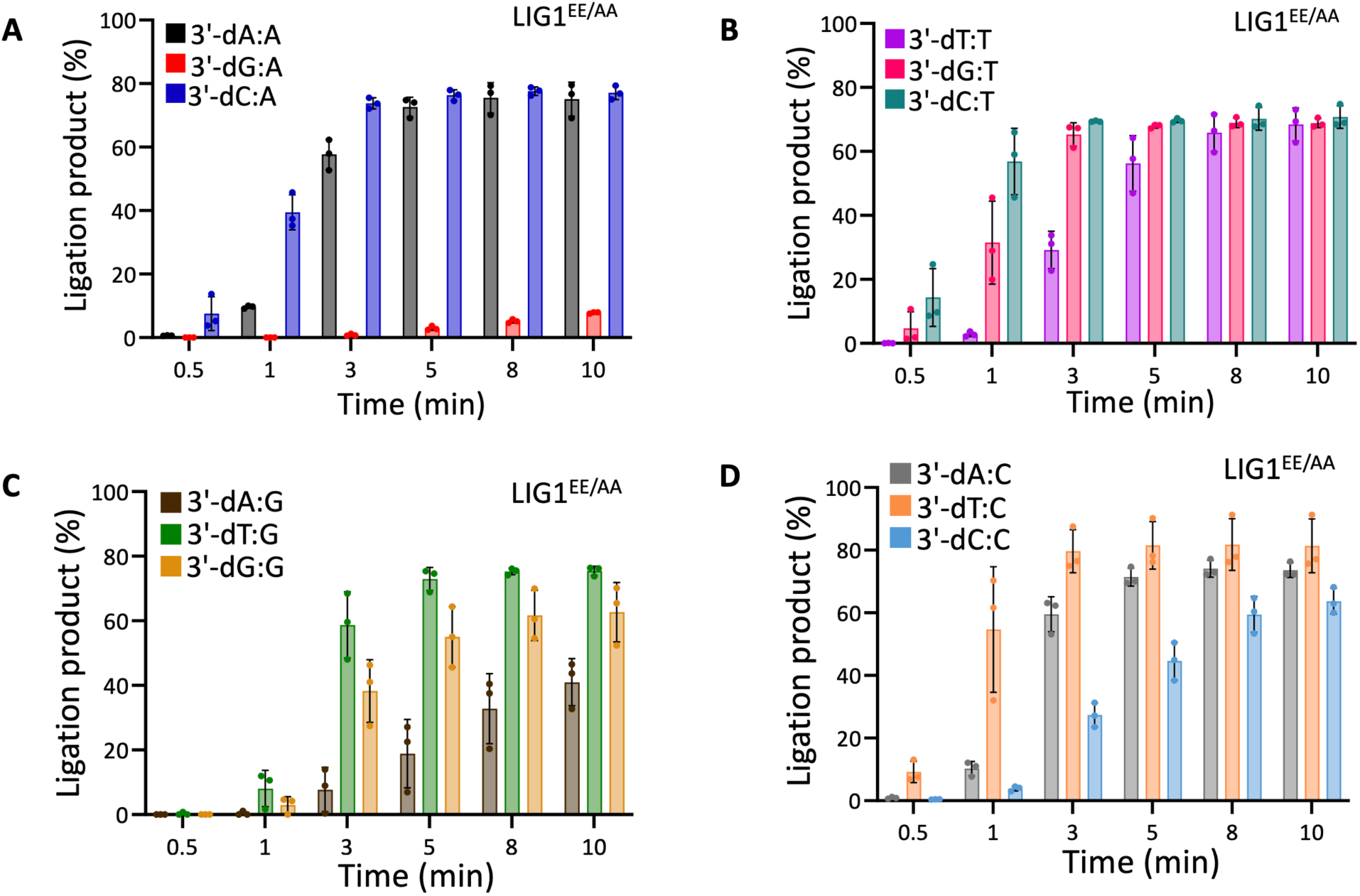
Ligation of nick DNA substrates containing template A, T, G and C mismatches by LIG1^EE/AA^ double mutant. (**A-D**) Graphs show time-dependent changes in the amounts of ligation products in the presence of template A (A), T (B), G (C), and C (D) mismatches by LIG1^EE/AA^ mutant. The data represent the average from three independent experiments ± SD.

**Supplementary Figure 7.**
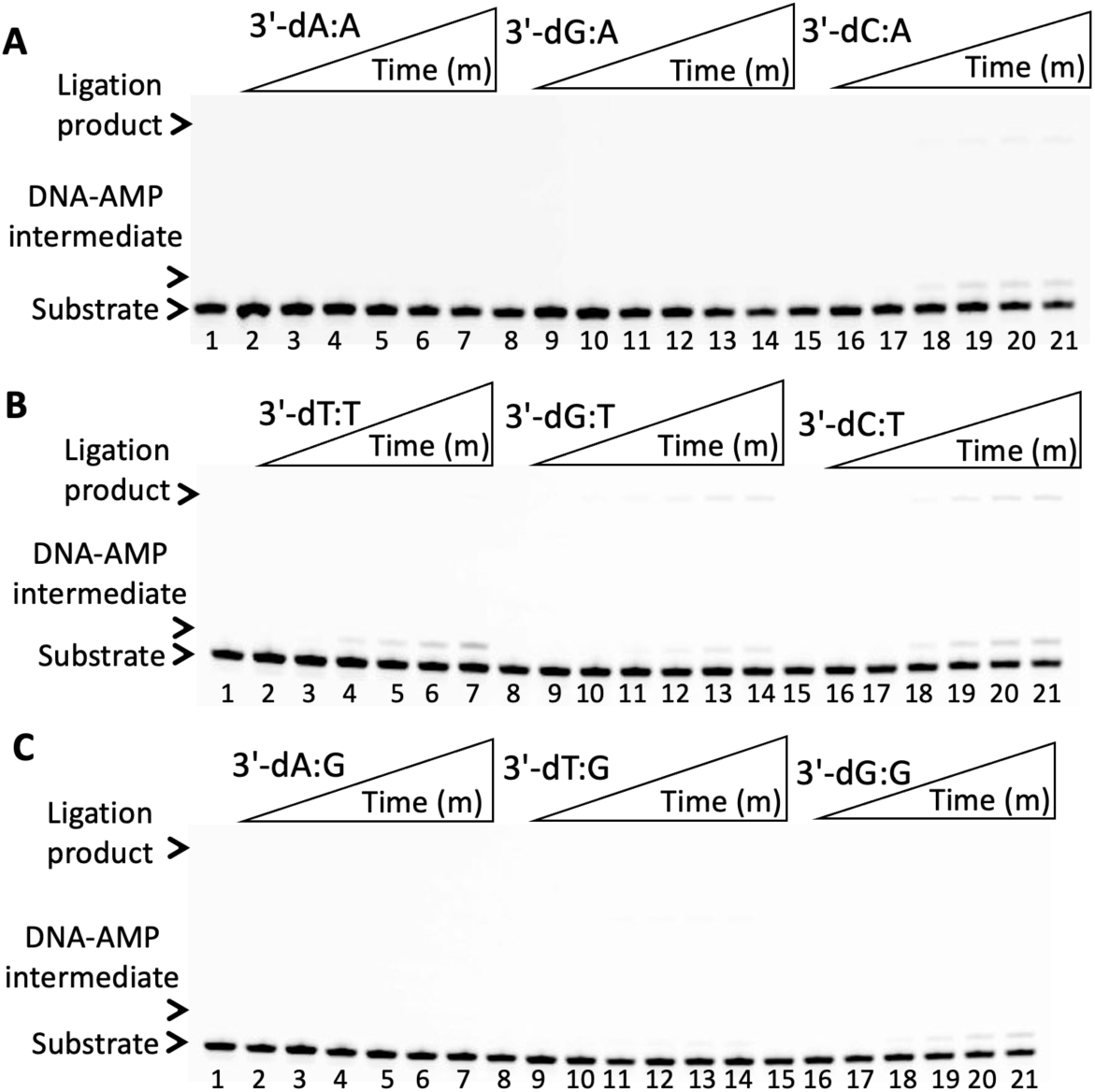
Ligation of nick DNA substrates containing template A, T, and G mismatches by LIG1 F635A single mutant. (**A-C**) Lanes 1, 8, and 15 are the negative enzyme controls of the nick DNA substrates containing template A, T, and G mismatches. Lanes 2-7, 9-14, and 16-21 are the ligation products in the presence of the nick DNA substrates with 3’-mismatches by LIG1 F635A mutant, and correspond to time points of 0.5, 1, 3, 5, 8, and 10 min. Graphs showing time-dependent changes in the amount of ligation products are presented in Figure 5.

**Supplementary Figure 8.**
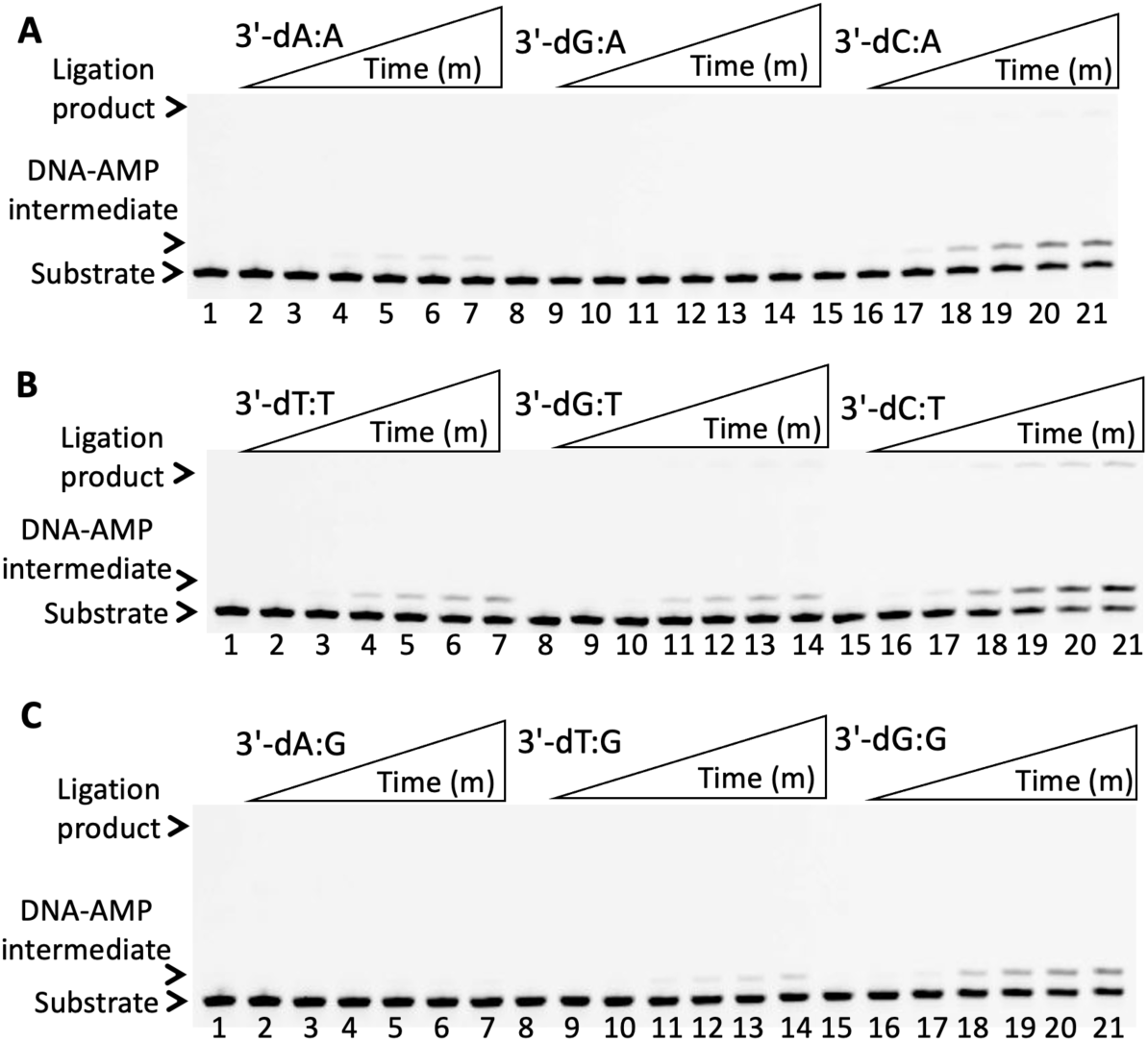
Ligation of nick DNA substrates containing template A, T, and G mismatches by LIG1 F635L single mutant. (**A-C**) Lanes 1, 8, and 15 are the negative enzyme controls of the nick DNA substrates containing template A, T, and G mismatches. Lanes 2-7, 9-14, and 16-21 are the ligation products in the presence of the nick DNA substrates with 3’-mismatches by LIG1 F635L mutant, and correspond to time points of 0.5, 1, 3, 5, 8, and 10 min. Graphs showing time-dependent changes in the amount of ligation products are presented in Figure 6.

**Supplementary Figure 9.**
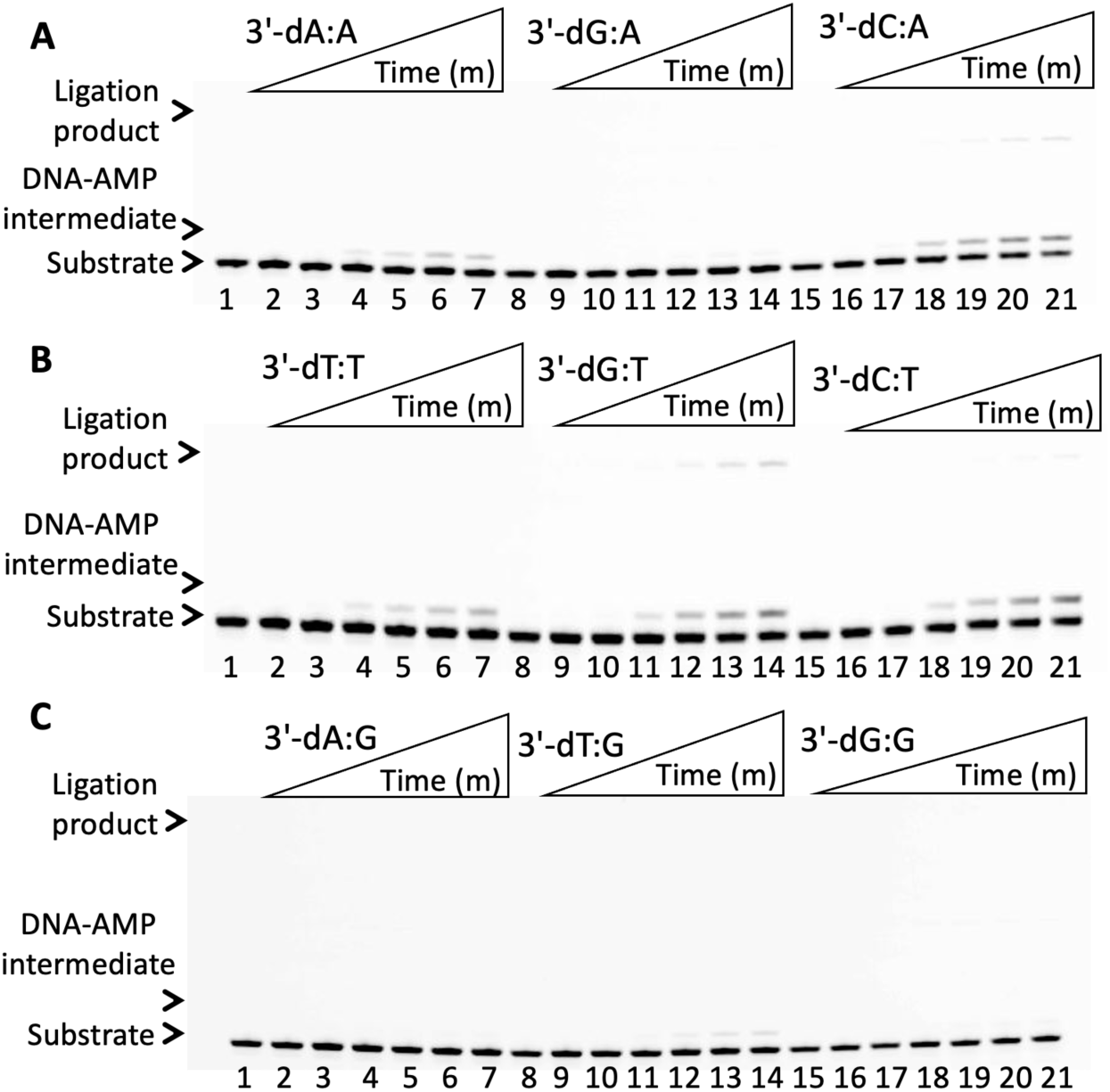
Ligation of nick DNA substrates containing template A, T, and G mismatches by LIG1 F872A single mutant. (**A-C**) Lanes 1, 8, and 15 are the negative enzyme controls of the nick DNA substrates containing template A, T, and G mismatches. Lanes 2-7, 9-14, and 16-21 are the ligation products in the presence of the nick DNA substrates with 3’-mismatches by F872A mutant, and correspond to time points of 0.5, 1, 3, 5, 8, and 10 min. Graphs showing time-dependent changes in the amount of ligation products are presented in Figure 7.

**Supplementary Figure 10.**
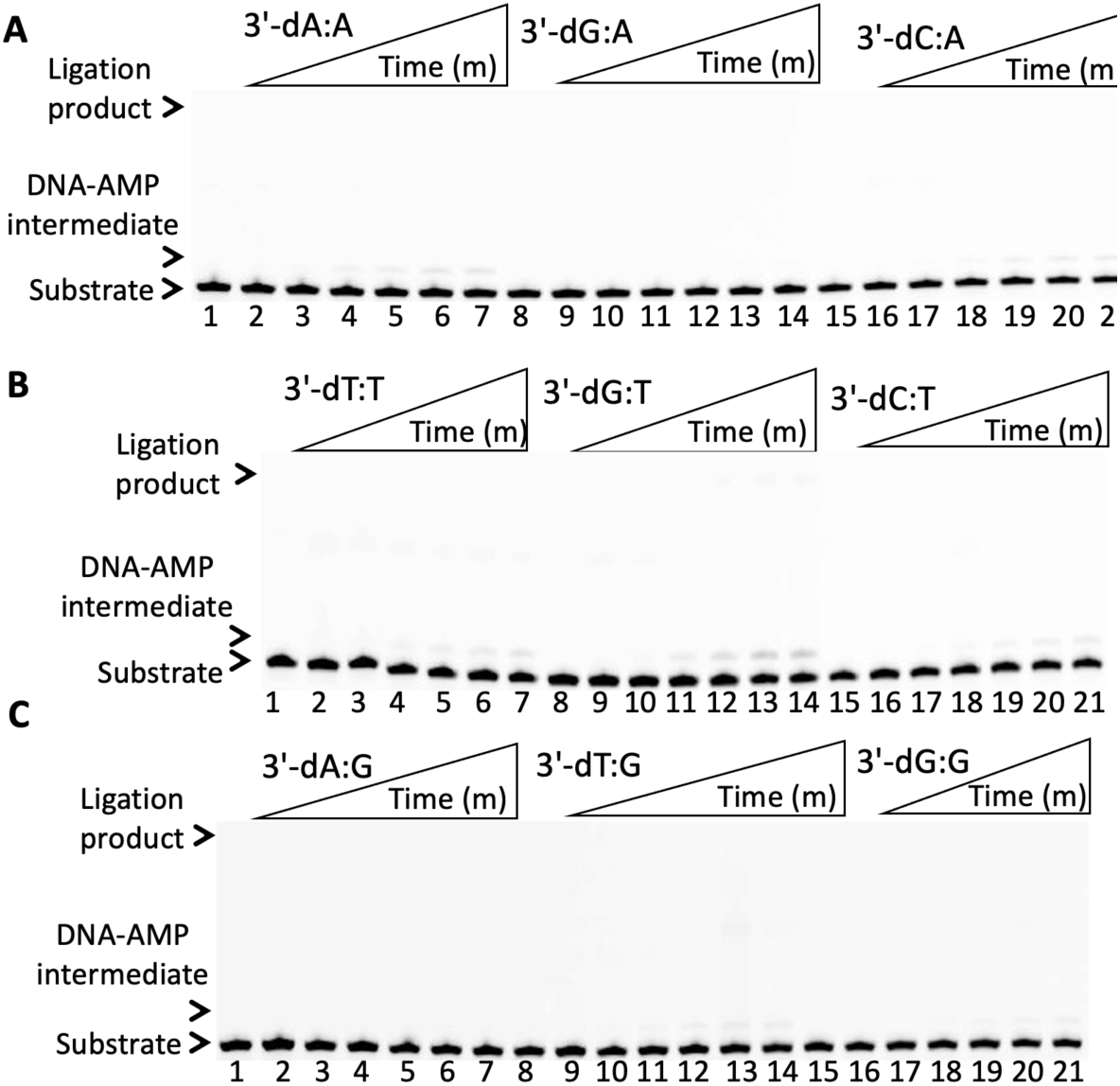
Ligation of nick DNA substrates containing template A, T, and G mismatches by LIG1 F872L single mutant. (**A-C**) Lanes 1, 8, and 15 are the negative enzyme controls of the nick DNA substrates containing template A, T, and G mismatches. Lanes 2-7, 9-14, and 16-21 are the ligation products in the presence of the nick DNA substrates with 3’-mismatches by F872L mutant, and correspond to time points of 0.5, 1, 3, 5, 8, and 10 min. Graphs showing time-dependent changes in the amount of ligation products are presented in Figure 8.

**Supplementary Figure 11.**
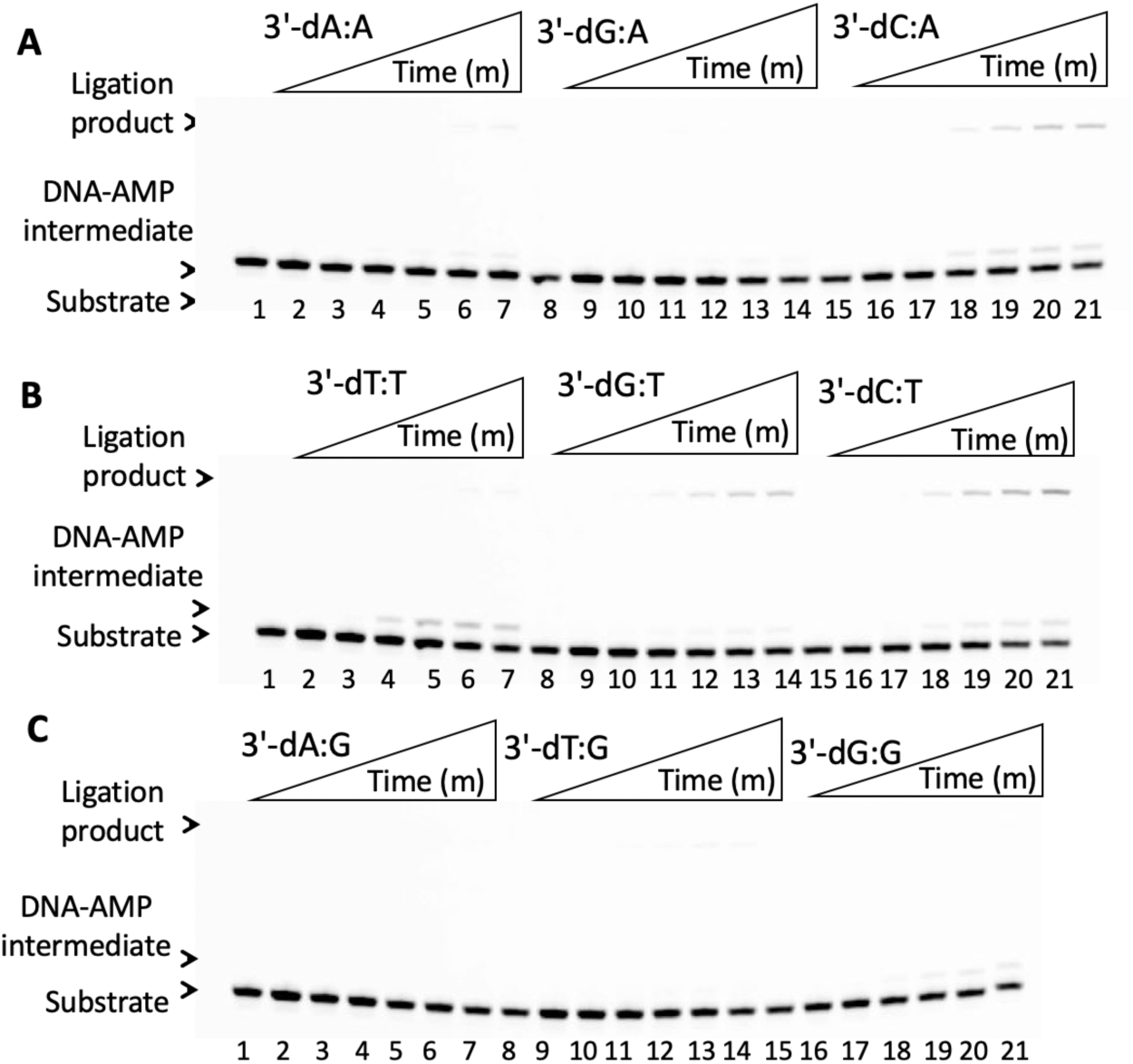
Ligation of nick DNA substrates containing template A, T, and G mismatches by LIG1^EE/AA^ F635A triple mutant. (**A-D**) Lanes 1, 8, and 15 are the negative enzyme controls of the nick DNA substrates containing template A, T, and G mismatches. Lanes 2-7, 9-14, and 16-21 are the ligation products in the presence of the nick DNA substrates with 3’-mismatches by LIG1^EE/AA^ F635A mutant, and correspond to time points of 0.5, 1, 3, 5, 8, and 10 min. Graphs showing time-dependent changes in the amount of ligation products are presented in Figure 9.

**Supplementary Figure 12.**
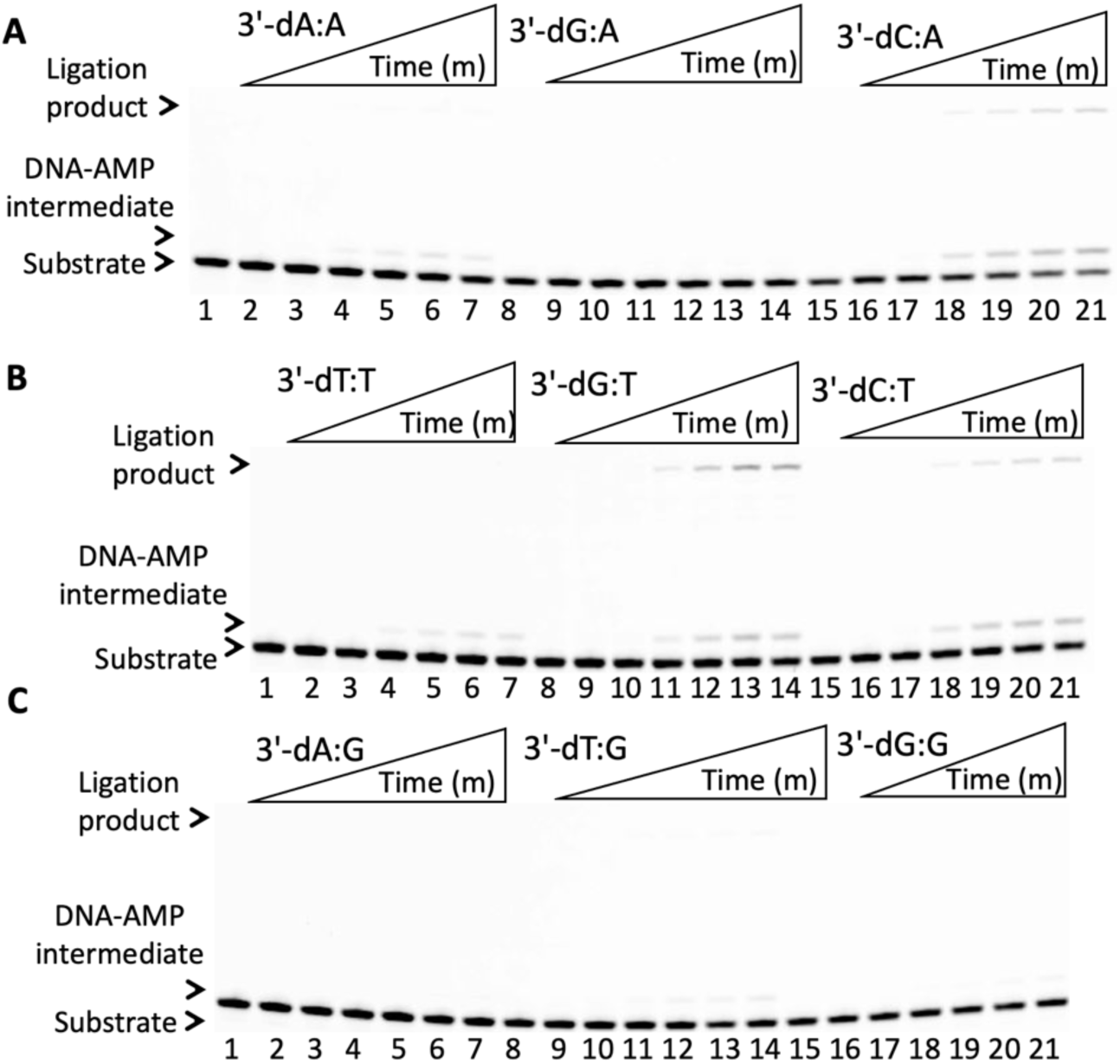
Ligation of nick DNA substrates containing template A, T, and G mismatches by LIG1^EE/AA^ F872A triple mutant. (**A-C**) Lanes 1, 8, and 15 are the negative enzyme controls of the nick DNA substrates containing template A, T, and G mismatches. Lanes 2-7, 9-14, and 16-21 are the ligation products in the presence of the nick DNA substrates with 3’-mismatches by LIG1^EE/AA^ F872A triple mutant, and correspond to time points of 0.5, 1, 3, 5, 8, and 10 min. Graphs showing time-dependent changes in the amount of ligation products are presented in Figure 10.

**Supplementary Figure 13.**
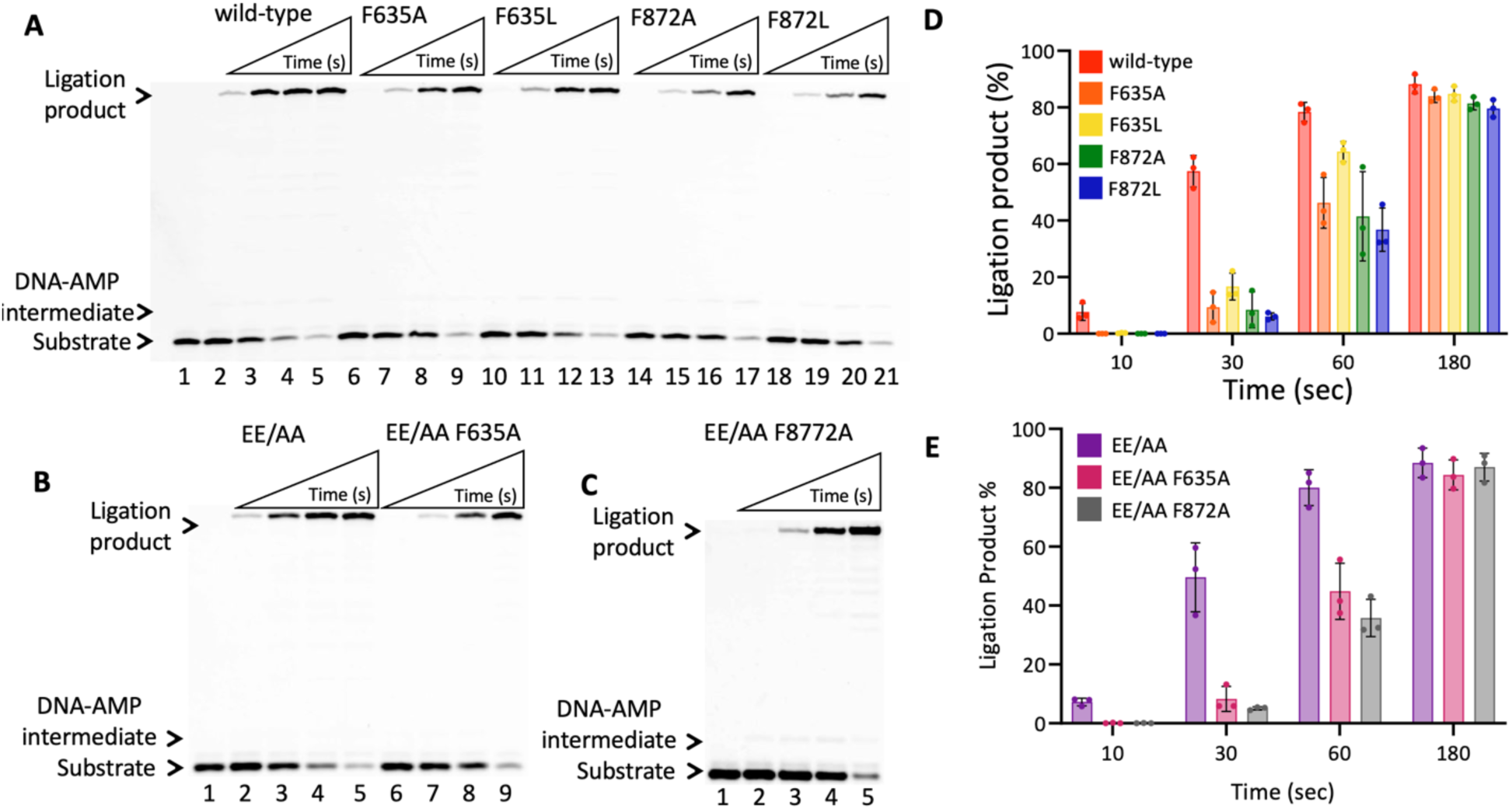
Ligation of nick DNA substrate with 3’-dG:C by LIG1 wild-type and seven mutants tested in this study. (**A**) Lane 1 is the negative enzyme control of the nick DNA substrate containing 3’-dG:C. Lanes 2-5, 6-9, 10-13, 14-17, and 18-21 are the ligation products by LIG1 wild-type, single mutants F635A, F635L, F872A, and F872L, and correspond to time points of 10, 30, 60, and 120 sec. (**B**) Lane 1 is the negative enzyme control of the nick DNA substrate containing 3’-dG:C. Lanes 2-5 and 6-9 are the ligation products by LIG1^EE/AA^ double and LIG1^EE/AA^ F635A triple mutants, and correspond to time points of 10, 30, 60, and 120 sec. (**C**) Lane 1 is the negative enzyme control of the nick DNA substrate containing 3’-dG:C. Lanes 2-5 are the ligation products by LIG1^EE/AA^ F872A triple mutant and correspond to time points of 10, 30, 60, and 120 seconds. (**D-E**) Graphs show time-dependent changes in the amounts of ligation products. The data represent the average from three independent experiments ± SD.

**Supplementary Figure 14.**
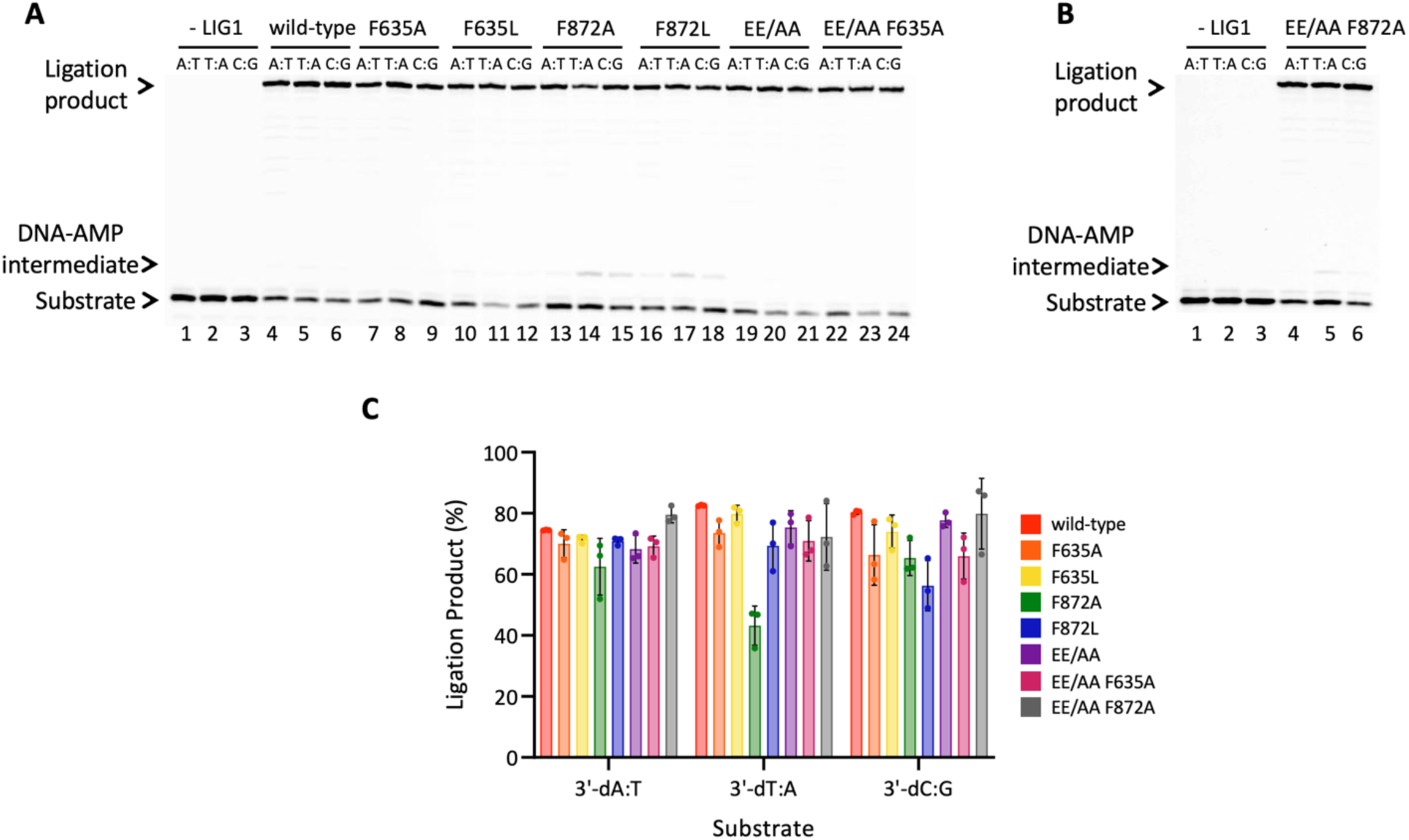
Ligation of nick DNA substrate with 3’-cognate base pairs A:T, T:A, and C:G by LIG1 wild-type and seven mutants tested in this study. (**A**) Lanes 1-3 are the negative enzyme controls of the nick DNA substrates containing 3’-preinserted dA:T, dT:A, and dC:G. Lanes 4-24 are the ligation products for 3 minutes by LIG1 wild-type, single mutants F635A, F635L, F872A, F872L, double mutant LIG1^EE/AA^, and triple mutant LIG1^EE/AA^ F635A. (**B**) Lanes 1-3 are the negative enzyme controls of the nick DNA substrates containing 3’-preinserted dA:T, dT:A, and dC:G. Lanes 4-6 are the ligation products for 3 minutes by triple mutant LIG1^EE/AA^ F872A. (**C**) Graph shows the amounts of ligation products for nick DNA substrates with 3’-dA:T, 3’-dT:A, and 3’-dC:G. The data represent the average from three independent experiments ± SD.

**Supplementary Figure 15.**
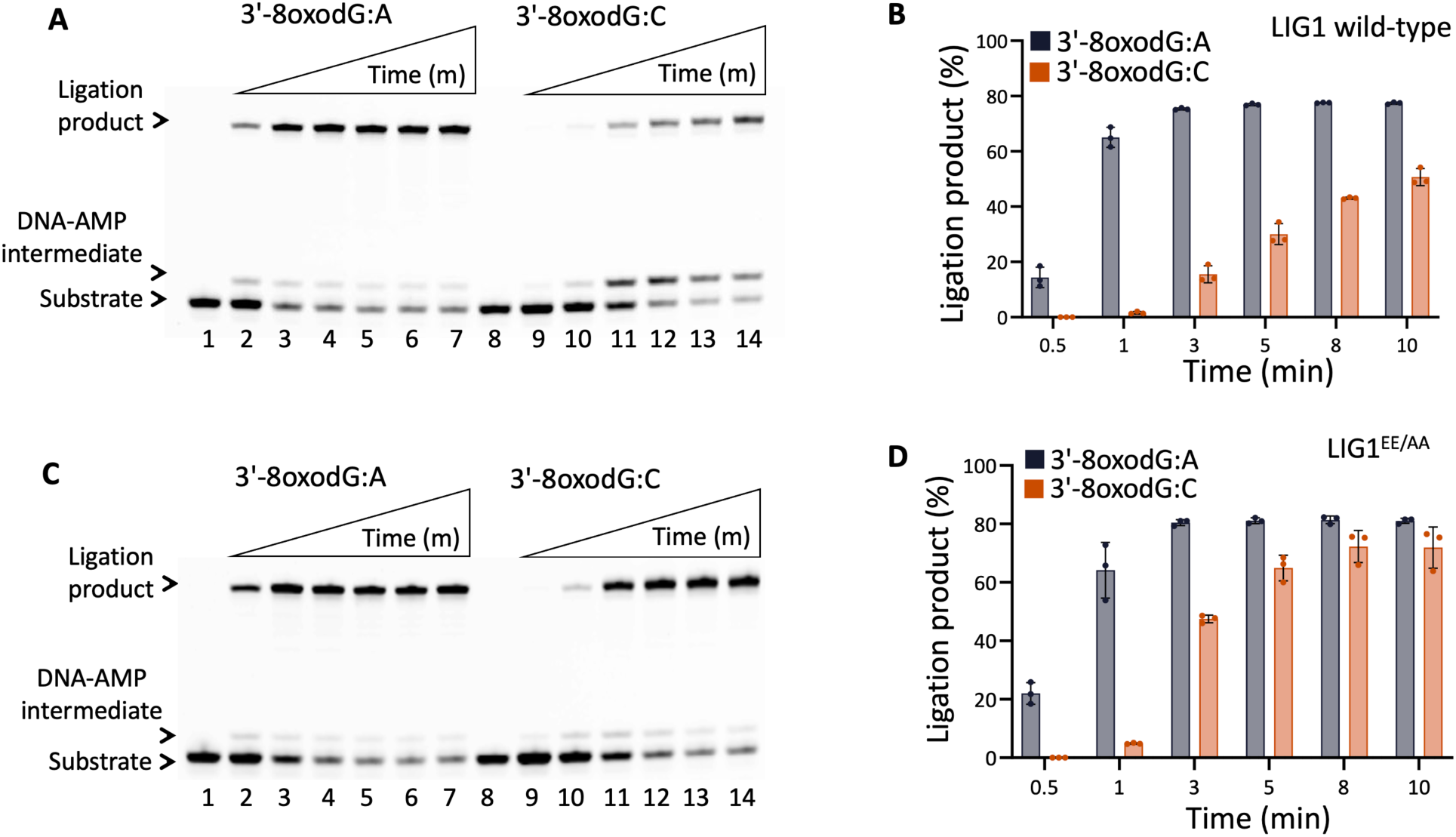
Ligation of nick DNA substrates containing 3’-8oxoG by wild-type LIG1 and LIG1^EE/AA^ mutant. (**A**) Lanes 1 and 8 are the negative enzyme controls of the nick DNA substrates containing 3’-8oxoG:A and 3’-8oxoG:C, respectively. Lanes 2-7 and 9-14 are the ligation products of nick DNA substrates with 3’-mismatches by wild-type LIG1, and correspond to time points of 0.5, 1, 3, 5, 8, and 10 min. (**B**) Graph shows the amounts of ligation products for nick DNA substrates with 3’-8oxoG:A and 3’-8oxoG:C. The data represent the average from three independent experiments ± SD. (**C**) Lanes 1 and 8 are the negative enzyme controls of the nick DNA substrates containing 3’-8oxoG:A and 3’-8oxoG:C, respectively. Lanes 2-7 and 9-14 are the ligation products of nick DNA substrates with 3’-mismatches by LIG1^EE/AA^ mutant, and correspond to time points of 0.5, 1, 3, 5, 8, and 10 min. (**D**) Graph shows the amounts of ligation products for nick DNA substrates with 3’-8oxoG:A and 3’-8oxoG:C. The data represent the average from three independent experiments ± SD.

**Supplementary Figure 16.**
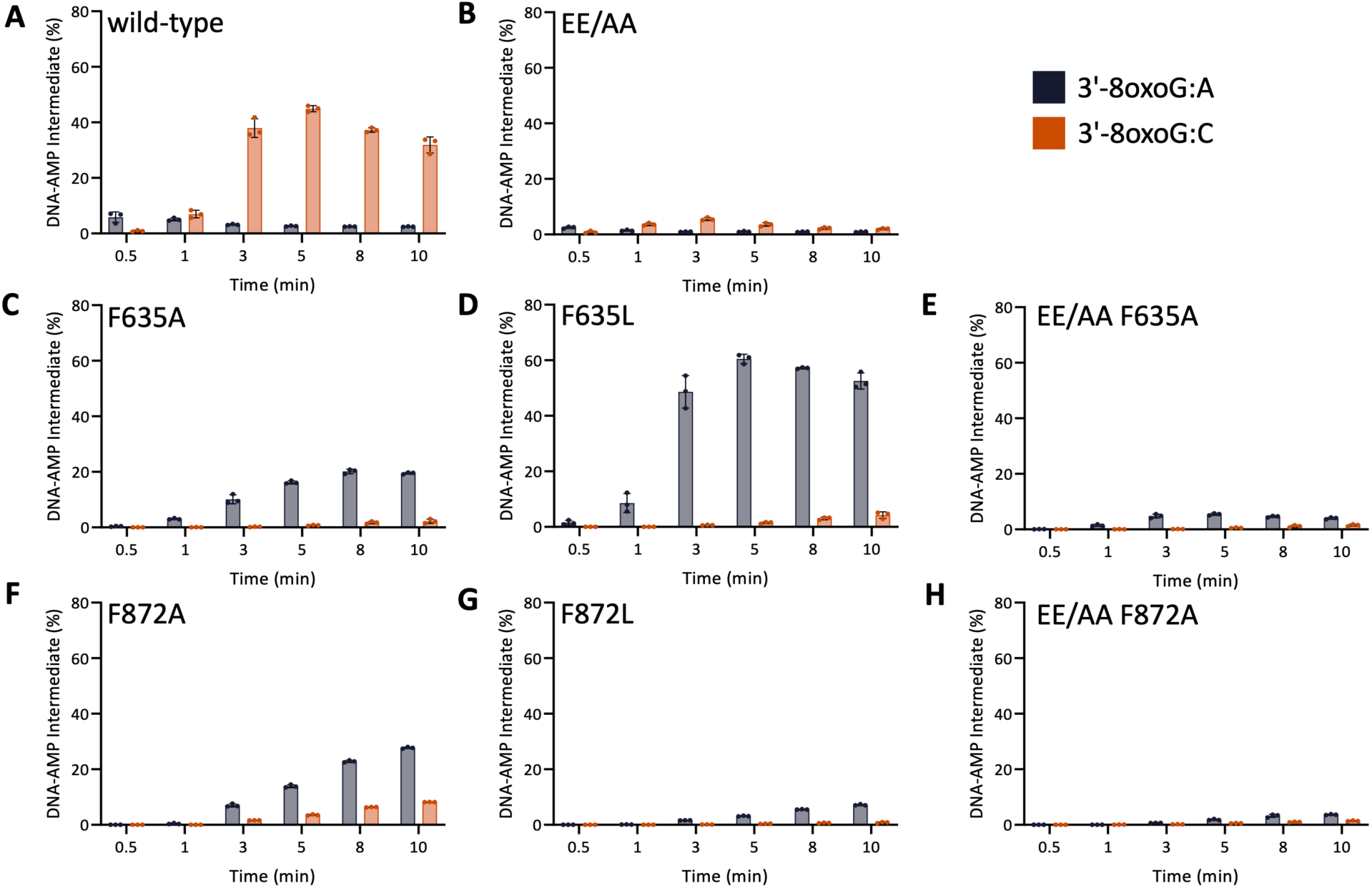
Comparison of DNA-AMP intermediates formed during the ligation of nick DNA substrates with 3’-8oxoG by LIG1 wild-type and seven mutants tested in this study. (**A-H**) Graphs show the amounts of DNA-AMP intermediates for nick DNA substrates with 3’-8oxoG by wild-type (A), LIG1^EE/AA^ (B), single mutants F635A (C), F635L (D), triple mutant LIG1^EE/AA^ F635A (E), single mutants F872A (F), F872L (G), and triple mutant LIG1^EE/AA^ F872A (H). The data represent the average from three independent experiments ± SD.

**Supplementary Figure 17.**
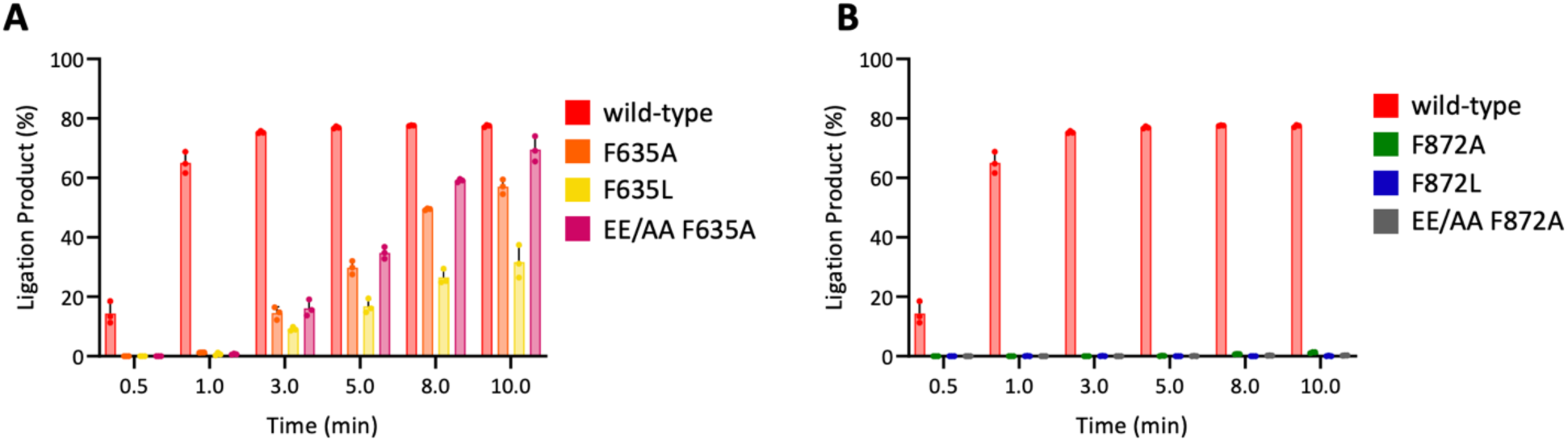
Comparison of ligation products for nick DNA substrates with 3’-8oxoG between LIG1 mutants tested in this study. (**A**) Comparison of the time-dependent ligation efficiency of the nick DNA substrate with 3’-8oxoG:A by LIG1 wild-type *versus* single mutants F635A and F635L as well as triple mutant LIG1^EE/AA^ F635A. (**B**) Comparison of the time-dependent ligation efficiency of the nick DNA substrate with 3’-8oxoG:A by LIG1 wild-type *versus* single mutants F872A and F872L as well as triple mutant LIG1^EE/AA^ F872A.

**Supplementary Figure 18.**
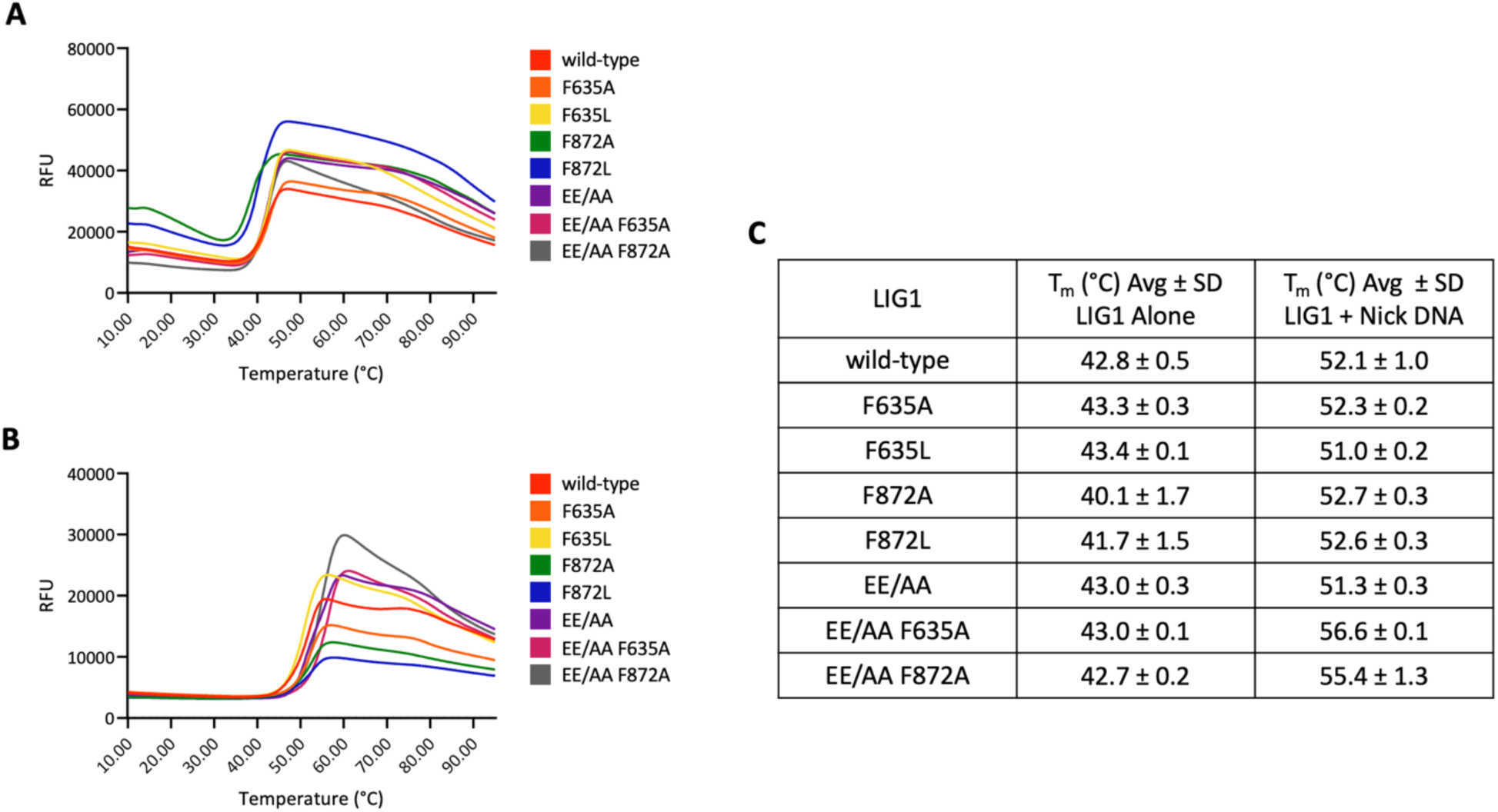
Thermal shift assay of LIG1 wild-type and seven mutants used in this study. (**A-B**) Plots show temperature-dependent change in SYPRO Orange fluorescence for LIG1 wild-type, single mutants F635A, F635L, F872A, F872L, LIG1^EE/AA^ mutant, triple-mutants LIG1^EE/AA^ F635A and F872A in the absence (**A**) and presence (**B**) of nick DNA. (**C**) Table shows the average T_m_ values ± SD for each LIG1 protein. T_m_ values were calculated manually using the inverse derivative plot of RFU values using the average of at least two biological repeats containing three technical repeats each.

**Supplementary Scheme 1.**
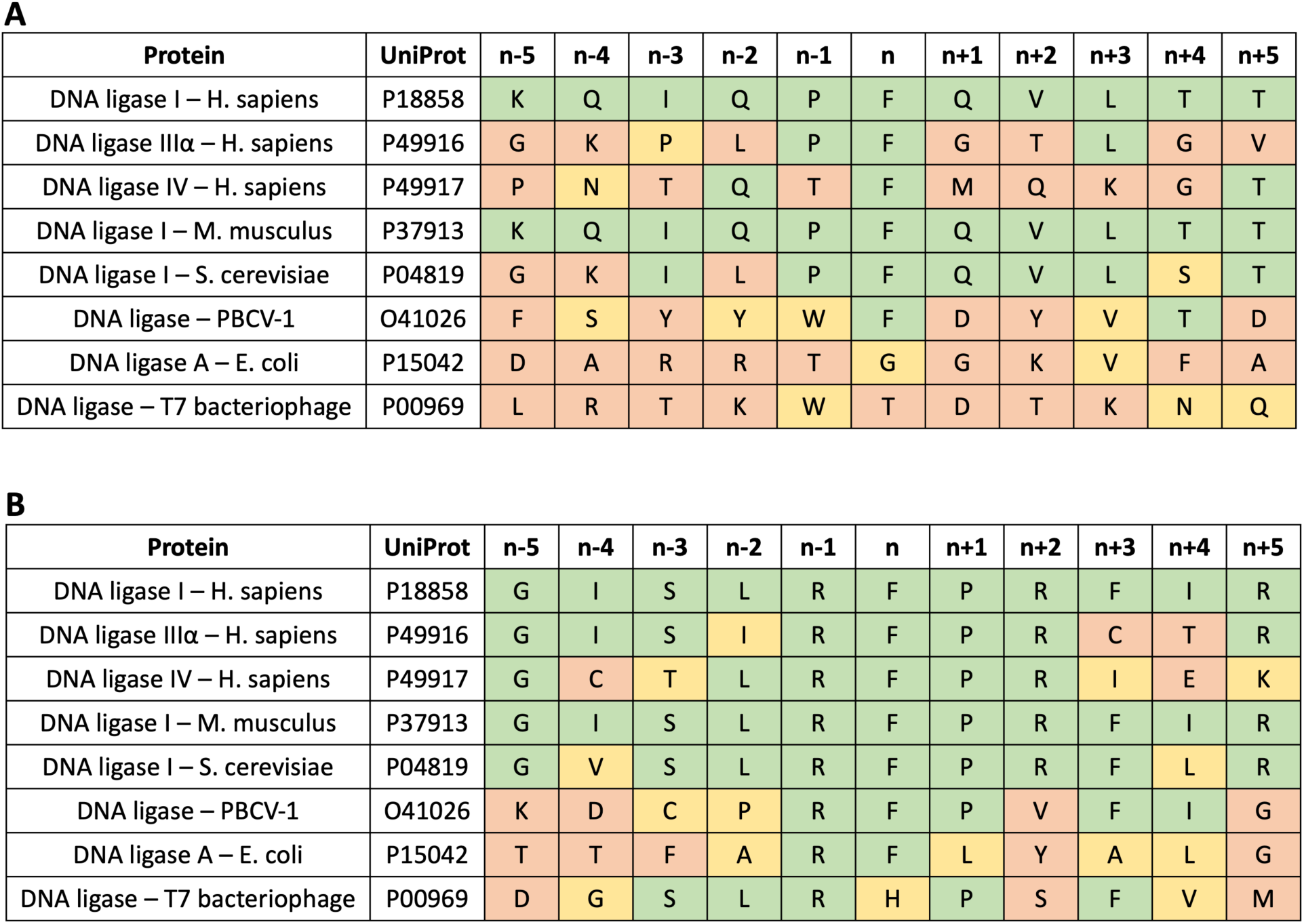
Sequence alignment of DNA ligases for active site residues F635 and F872. (**A**) The amino acid sequence alignment for the region corresponding to 629-640 of LIG1 demonstrates the conservation of the phenylalanine residue at F635. (**B**) The amino acid sequence alignment for the region corresponding to 867-877 of LIG1 demonstrates the conservation of a phenylalanine residue at F872. In both, green represents an exact amino acid match, yellow represents an amino acid class match (polar, nonpolar, positively charged, or negatively charged), and red represents no match.

**Supplementary Scheme 2.**
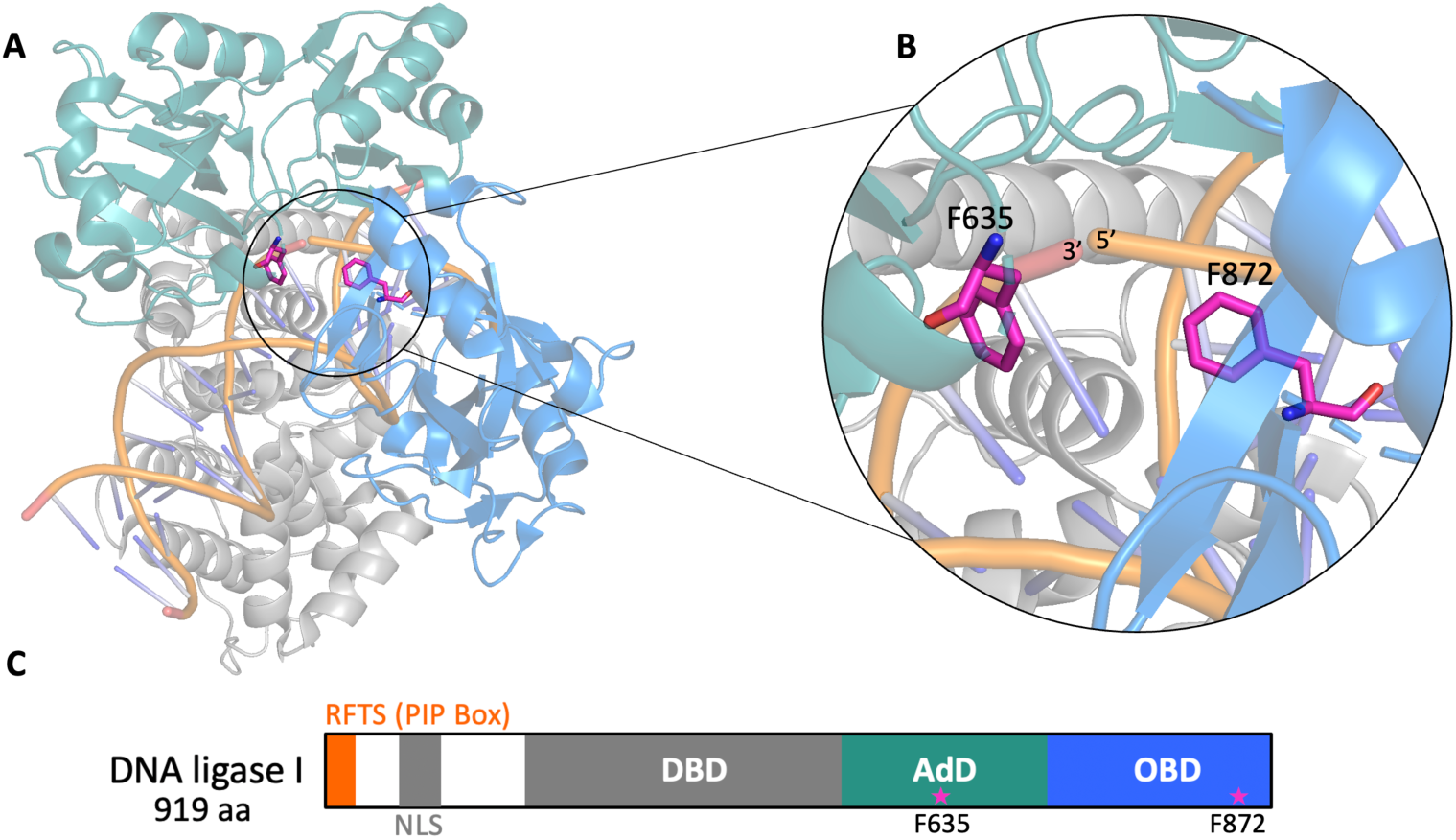
Domain organization of human LIG1. (**A**) Structure of the C-terminal globular domains of LIG1 (PDB: 6P0C). (**B**) Zoomed view of the LIG1 active site. F635 and F872 are shown as sticks in magenta. The 3’- and 5’-ends of the nick are labeled. (**C**) Domain organization map of human LIG1 protein (1-919 amino acids). The DNA binding domain (DBD), the adenylation domain (AdD), and the oligonucleotide/oligosaccharide binding domain (OBD) are colored in grey, teal, and blue, respectively. RFTS refers to the replication factor targeting sequence where the PIP box is located. NLS stands for nuclear localization signal. The amino acid residues are marked where each domain starts and ends. The position of F635 and F872 are shown as magenta stars.

**Supplementary Scheme 3.**
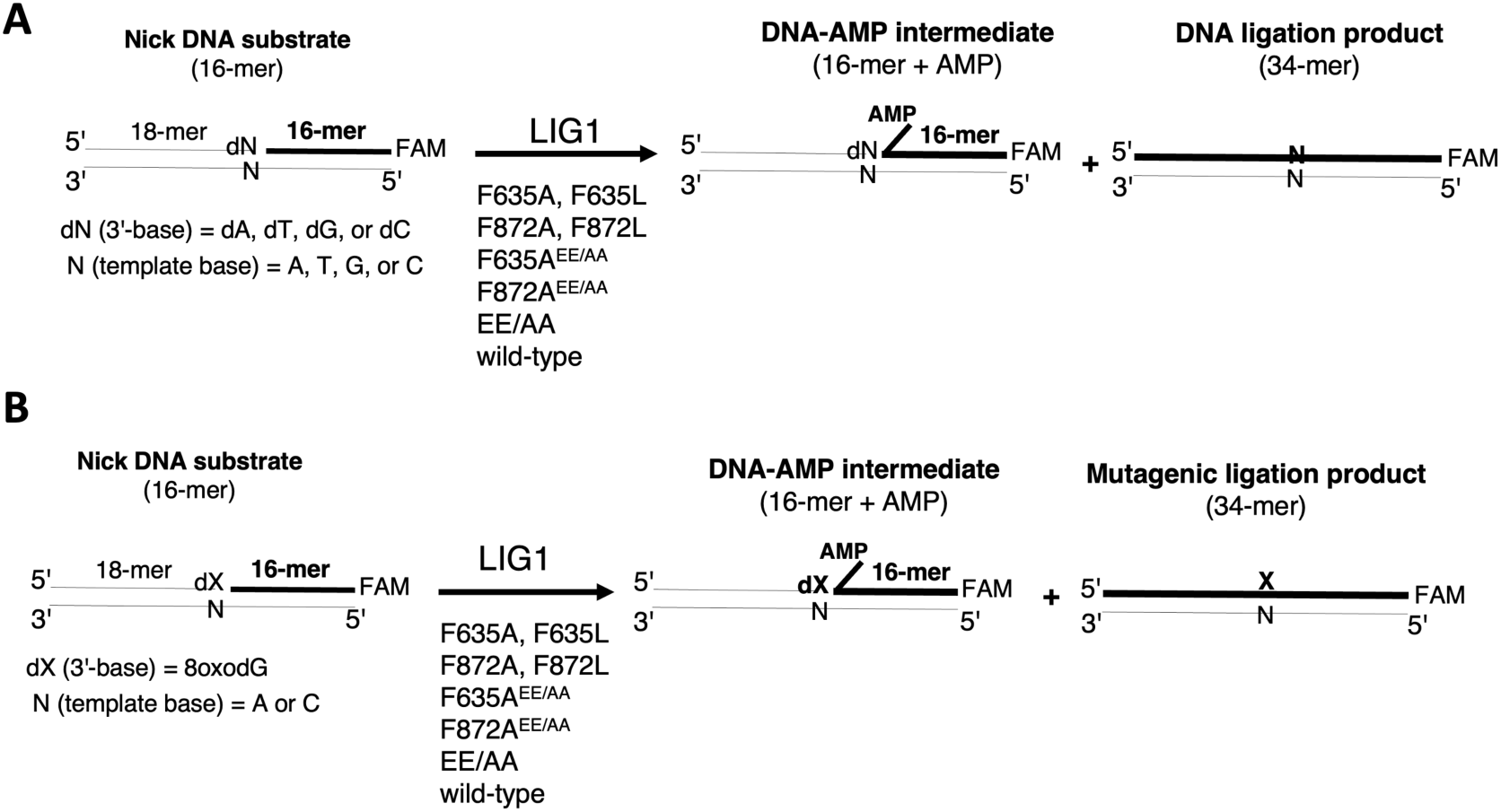
Illustrations of DNA ligation assays used in this study. Ligation assays were used to evaluate the substrate specificity of LIG1 wild-type and active site mutants for the nick DNA substrates including 3’-mismatches (**A**) and 3’-8oxoG (**B**). Reaction products include nick sealing/ligation product and DNA-AMP intermediates with 5’-adenylate (AMP).

**Supplementary Table 1.**
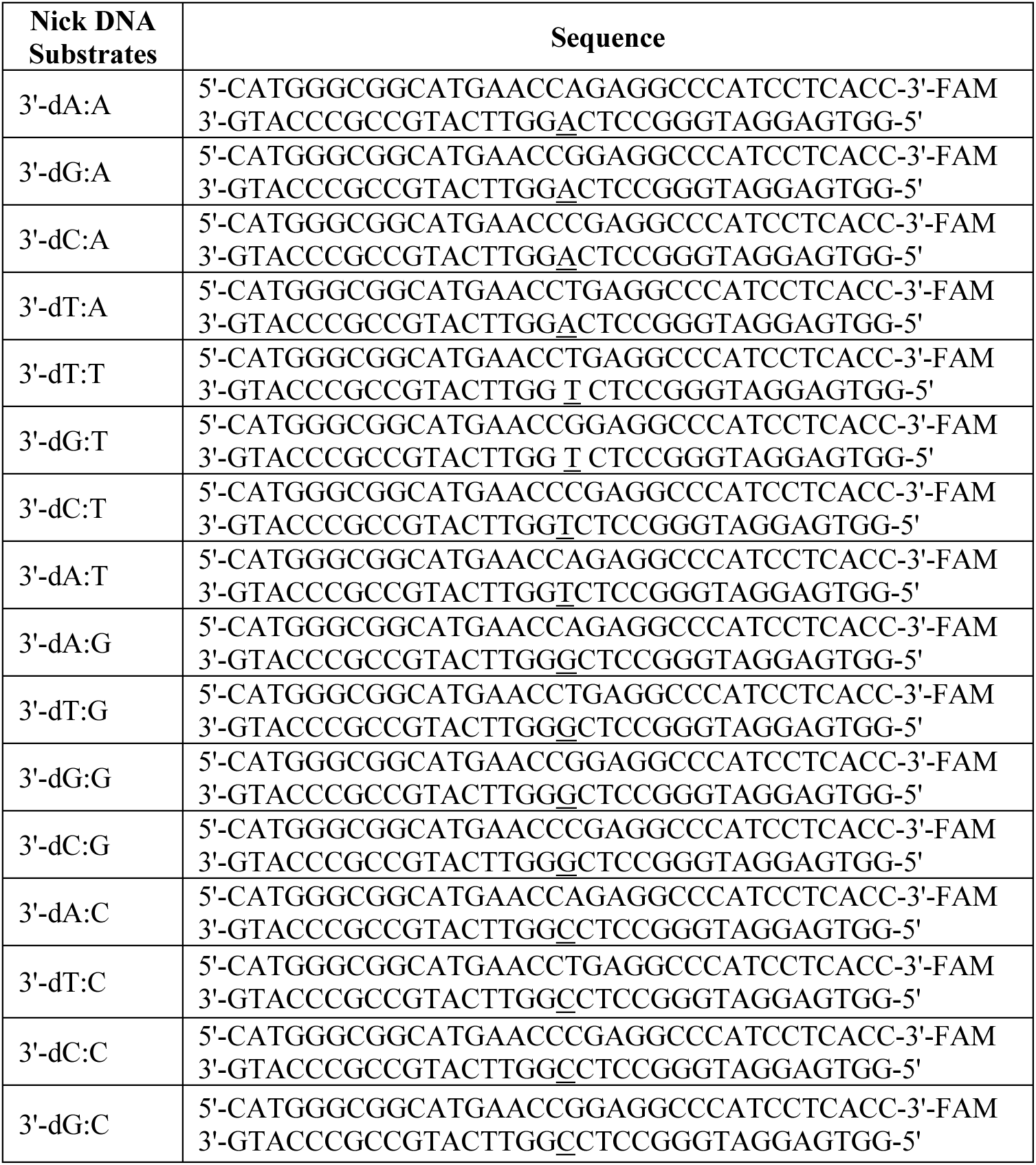
Nick DNA substrates containing 3’-preinserted mismatches. Nick DNA substrates with preinserted 3’-dA, dT, dG, dC opposite template base A, T, G, or C were used in the ligation assays (Supplementary Scheme 3A). FAM denotes a fluorescent tag and is located at 3’-end of DNA substrates. The base at the template position is underlined.

**Supplementary Table 2.**
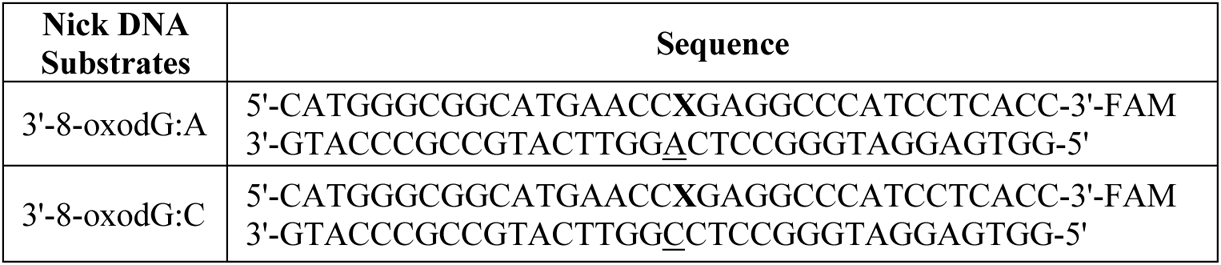
Nick DNA substrates containing 3’-8oxodG. Nick DNA substrates with 3’-8oxodG opposite template base A or C were used in the ligation assays (Supplementary Scheme 3B). FAM denotes a fluorescent tag and is located at 3’-end of DNA substrates. X represent 8-oxodG and the base at the template position is underlined.

**Supplementary Table 3:**
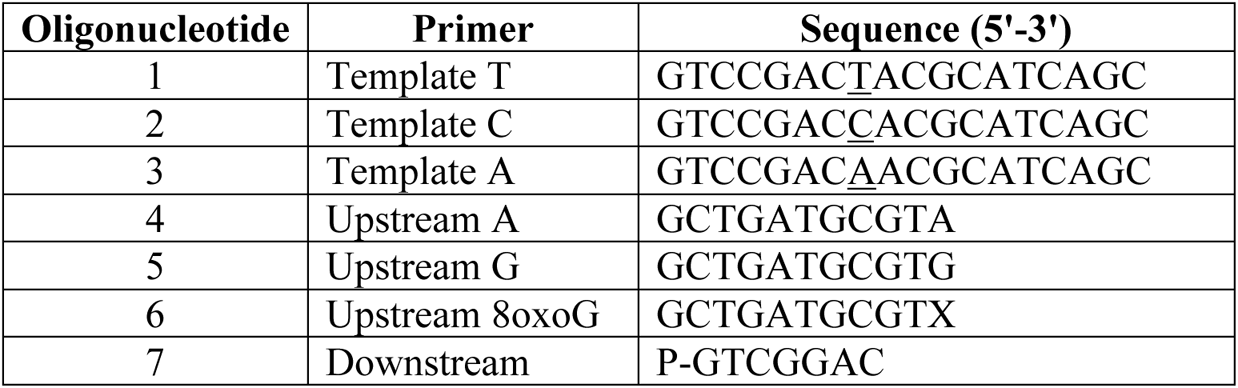
Oligonucleotides used for LIG1 crystallization. Oligonucleotides 1, 4, and 7 were used to prepare the nick DNA substrate with 3’-A:T. Oligonucleotides 2, 4, and 7 were used to prepare the nick DNA substrate with 3’-A:C. Oligonucleotides 1, 5, and 7 were used to prepare the nick DNA substrate with 3’-G:T. Oligonucleotides 3, 6, and 7 were used to prepare the nick DNA substrate with 3’-8oxoG:A. The base at the template position is underlined. X represents 8oxoG. P denotes a phosphate group at the 5’-end.

**Supplementary Table 4.**
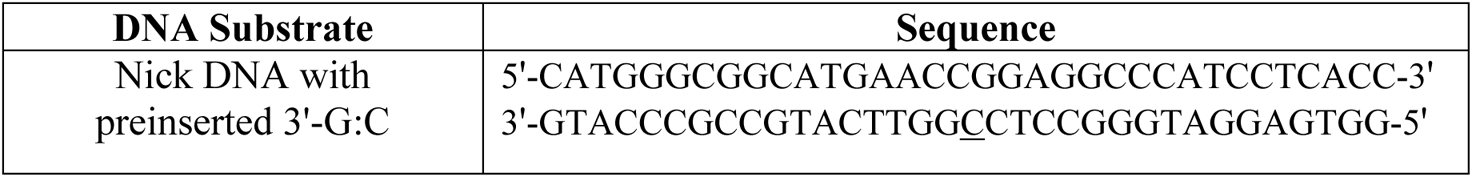
Nick DNA substrate used in thermal shift assays. Nick DNA substrate with 3’-dG:C was used in the thermal shift assays. The base at the template position is underlined.

**Supplementary Table 5.** Crystallization conditions used for LIG1 structures of the present study. Reservoir solutions used in hanging drop crystallization experiments to grow diffraction quality crystals for data collection.

| Protein/DNA | Crystallization Conditions |
| --- | --- |
| LIG1<sup>EE/AA</sup> F635A/A:T | 100 mM MES pH 6.9, 20% PEG3350 (w/v), 100 mM Lithium Acetate |
| LIG1<sup>EE/AA</sup> F635A/A:C | 100 mM MES pH 6.3, 24% PEG3350 (w/v), 25 mM Lithium Acetate |
| LIG1<sup>EE/AA</sup> F635A/G:T | 100 mM MES pH 6.7, 26% PEG3350 (w/v), 100 mM Lithium Acetate |
| LIG1<sup>EE/AA</sup> F635A/8oxoG:A | 100 mM MES pH 6.4, 20% PEG3350 (w/v), 100 mM Lithium Acetate |
| LIG1<sup>EE/AA</sup> F872A/A:T | 100 mM MES pH 6.7, 16% PEG3350 (w/v), 100 mM Lithium Acetate |
| LIG1<sup>EE/AA</sup> F872A/A:C | 100 mM MES pH 5.7, 20% PEG3350 (w/v), 100 mM Lithium Acetate |
| LIG1<sup>EE/AA</sup> F872A/8oxoG:A | 100 mM MES pH 6.4, 24% PEG3350 (w/v), 100 mM Lithium Acetate |

## Notes

### Competing Interest Statement

The authors have declared no competing interest.

